# A unique conformational escape reaction of HIV-1 against an allosteric integrase inhibitor

**DOI:** 10.1101/836718

**Authors:** Tomofumi Nakamura, Teruya Nakamura, Masayuki Amano, Toshikazu Miyakawa, Yuriko Yamagata, Masao Matsuoka, Hirotomo Nakata

## Abstract

HIV-1 integrase (IN) contributes to HIV-1 RNA binding, which is required for viral maturation. Non-catalytic site integrase inhibitors (NCINIs) have been developed as allosteric IN inhibitors, which perform anti-HIV-1 activity by disrupting IN multimerization. Here, we show that IN undergoes a novel conformational alteration to escape from NCINIs. We observed that NCINI-resistant HIV-1 variants have accumulated amino acid (AA) mutations in the IN-encoding region. We employed HPLC and thermal stability assays to show that the AA mutations affect the folding and dimerization interface of the IN catalytic core domains, resulting in severely decreased multimerization of full-length IN proteins (IN under-multimerization). The under-multimerization of IN was finally restored by HIV-1 RNA in the viral particles. Our study demonstrates that HIV-1 countervails NCINIs by IN under-multimerization as a novel escape mechanism. Our findings provide information on the understanding of IN multimerization and influence the development of unique anti-HIV-1 strategies.

## Introduction

Human immunodeficiency virus type 1 (HIV-1) infection has been a chronic infectious disease. Patients diagnosed with HIV are treated with anti-retroviral therapy (ART), which basically consists of more than two anti-HIV-1 drugs. The life expectancy of HIV-1-positive patients with ART has been prolonged to the almost same extent as that of uninfected individuals (***May et al., 2014***; ***Trickey et al., 2017***). Despite the benefits of ART, HIV-1 often acquires drug-resistant mutations (***Huldrych et al., 2018***), resulting in treatment failure. High adherence to ART is required to sustain viral suppression in HIV-1 clinical treatment (***Kanters et al., 2017***). Recently, the FDA has approved certain anti-HIV-1 drugs which can be easily taken once-daily in the form of a single tablet, greatly improving ART adherence (***Flexner et al., 2010***). Long-acting formulations of anti-HIV-1 drugs including a newer drug delivery system could be also useful in ART adherence and HIV-1 treatment. Moreover, HIV-1 prevention, pre-exposure prophylaxis (PrEP) by such long-acting formulations of anti-HIV-1 drugs, is currently being investigated in clinical trials (***Gulick et al., 2019***). On the other hand, studying the mechanism of HIV-1 resistance to anti-HIV-1 drugs would lead to the development of novel drugs with increased efficiency and a greater genetic barrier to resistance, resulting in more effective ART.

Four FDA-approved integrase strand transfer inhibitors (INSTIs), raltegravir (RAL), elvitegravir (ELG), dolutegravir (DTG), and bictegravir (BTG) which target the active site of IN are more potent and well tolerated than other classes of anti-HIV-1 drugs due to the lack of homologous human proteins, allowing these INSTIs to have been widely used for clinical HIV-1 treatment. The World Health Organization recommends INSTIs, especially DTG, for clinical HIV-1 treatment according to the December 2018 Interim guidelines. A single tablet containing only two anti-HIV-1 drugs (DTG and rilpivirine (RPV) - second-generation non-nucleoside reverse transcriptase inhibitor (NNRTIs)) was approved by the FDA in 2018 as the first maintenance therapy for HIV-1 infection (***Dowers et al., 2018***). Therefore, the development of more potent anti-HIV-1 inhibitor targets for IN would be useful for HIV-1 treatment.

The NCINIs were developed as novel integrase inhibitors interfering with IN binding to lens epithelium-derived growth factor p75 (LEDGF/p75) (***Christ et al., 2010***; ***Christ et al., 2012***) which plays an important role in tethering IN to the host genome and stabilizing functional IN multimerization (***Cherepanov et al., 2003***; ***Hare et al., 2009***). It was reported that certain more potent NCINIs induce over-multimerization of IN as the main anti-HIV-1 activity (***Tsiang et al., 2012***; ***Rouzic et al., 2013***; ***Sharma et al., 2014***), leading to the production of immature viruses in which the ribonucleoprotein (RNP) is translocated from the capsid core during HIV-1 maturation (***Desimmie et al., 2013***; ***Jurado et al., 2013***). Moreover, the immature and non-infectious HIV-1 produced by NCINIs resulted from loss of the interaction between over-multimerized IN and HIV-1 RNA during HIV-1 maturation (***Kessl et al., 2016***). However, the process of IN multimerization with HIV-1 RNA during HIV-1 maturation remain to be fully elucidated. In the present study, we acquired HIV-1 variants which are resistant to NCINIs disrupting IN multimerization, and attempted to examine the mechanism underlying the mutations which could provide details of IN multimerization during HIV-1 maturation. The study of drug-resistant mutations with respect to such allosteric inhibitors may help to elucidate how HIV-1 proteins undergo functional multimerization, which may aid in the development of novel anti-HIV-1 drugs and unique treatment strategies.

## Results

### Emergence of NCINI-related resistance mutations

NCINIs suppress replication of HIV-1 by inducing aberrant IN multimerization (IN over-multimerization) (***Kessl et al., 2012***; ***Tsiang et al., 2012***;***Jurado et al., 2013***; ***Rouzic et al., 2013***; ***Desimmie et al., 2013***). We selected HIV-1_NL4-3_ variants resistant to the three NCINIs (NCINI-1, -2, and -3) reported previously (***Christ et al., 2012***) by selection experiments (Figure 1 and Figure 1—figure supplement 1). As the concentration of the NCINIs up to 10 µM gradually increased, HIV-1 acquired several AA mutations, such as A128T and K173Q at passage 15 (P15) in the catalytic core domain (CCD), and N254K at passage 20 (P20) in the C-terminal domain (CTD) of IN. Four mutations, A128T, H171Q, K173Q, and N254K, accumulated during passage 26 (P26) in the full-length IN region upon treatment with NCINI-3 at 10 µM (Figure 1). A128T and G272G mutations were acquired in the IN region upon treatment with NCINI-1 at passage 18, and V77S and A128T mutations were acquired upon treatment with NCINI-2 at passage 12 (Figure 1).

**Figure 1.**
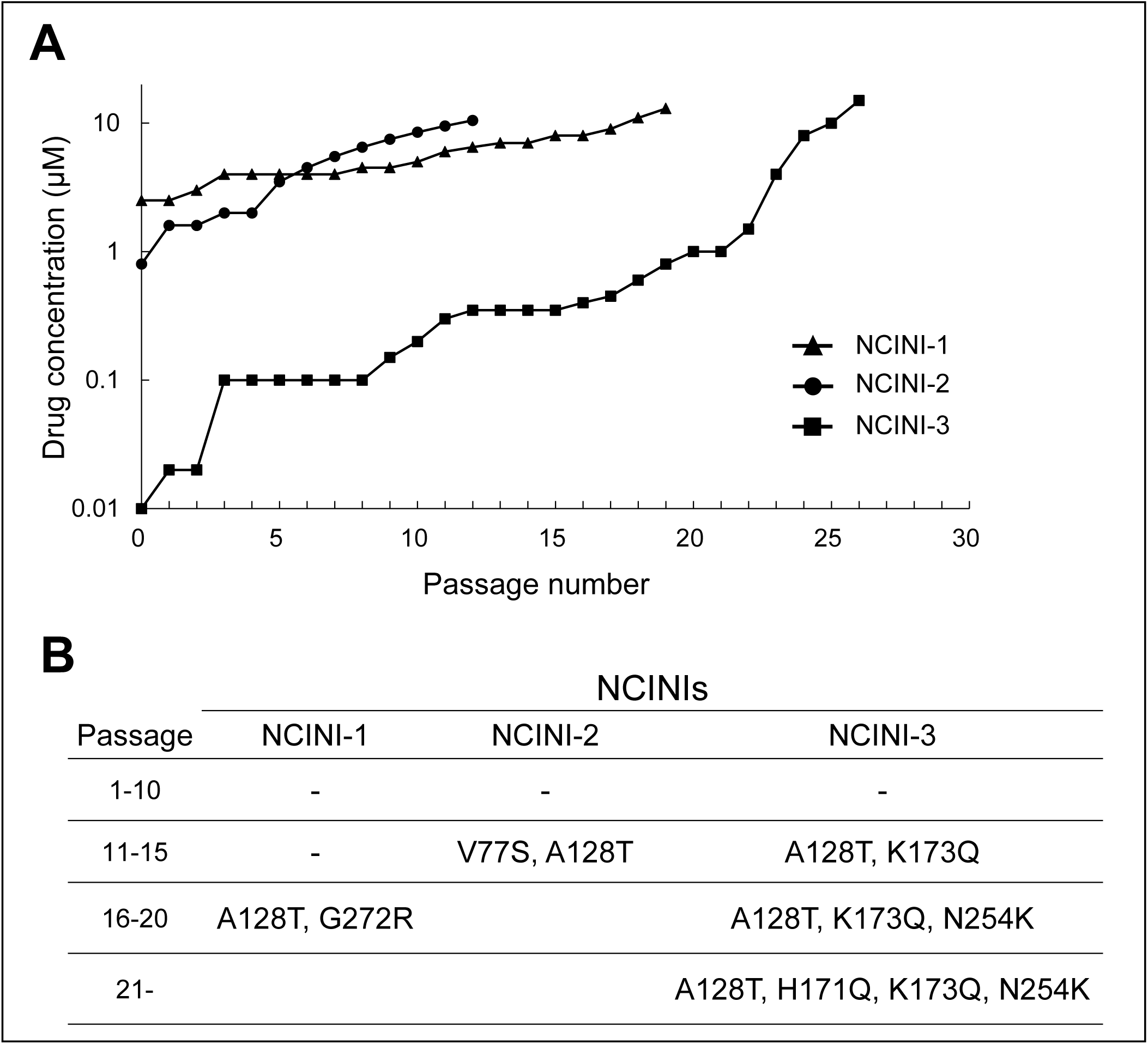
Emergence of NCINI-related resistance mutations in IN region. (A) HIV-1_NL4-3_ propagated in MT-4 cells in the presence of increasing concentrations of NCINI-1 (▴), NCINI-2 (●), or NCINI-3 (▪), and the resistant viral selection to the NCINIs continues up to 10 µM. (B) NCINI-related resistance mutations in the IN region are shown at following passage ranges 1-10, 11-15, 16-20, and over 21. These mutations at each passage are identified from cellular DNA in the infected MT-4 cells with the NCINI-related resistant HIV-1.

EC_50s_ of the drugs (NCINI-1, -2, -3, BI224436 (***Fenwick et al., 2014***), RAL, and darunavir (DRV)) against wild type HIV-1_NL4-3_ and recombinant HIV-1_NL4-3_ clones (HIV-1_NL4-3_-IN^A128T^, -IN^H171Q^, -IN^K173Q^, -IN^N254K^, -IN^P15^, -IN^P20^, and -IN^P26^, respectively), and fold changes (EC_50_ ratios against each HIV-1 clone to wild type HIV-1) are shown in Table 1 and Table 1—table supplement 1. Accumulation of these mutations gradually gave rise to NCINI-resistant HIV-1; however, none of the clones carrying NCINI-related resistance mutations exhibited cross resistance to clinically used drugs, such as RAL and DRV.

**Table 1.**
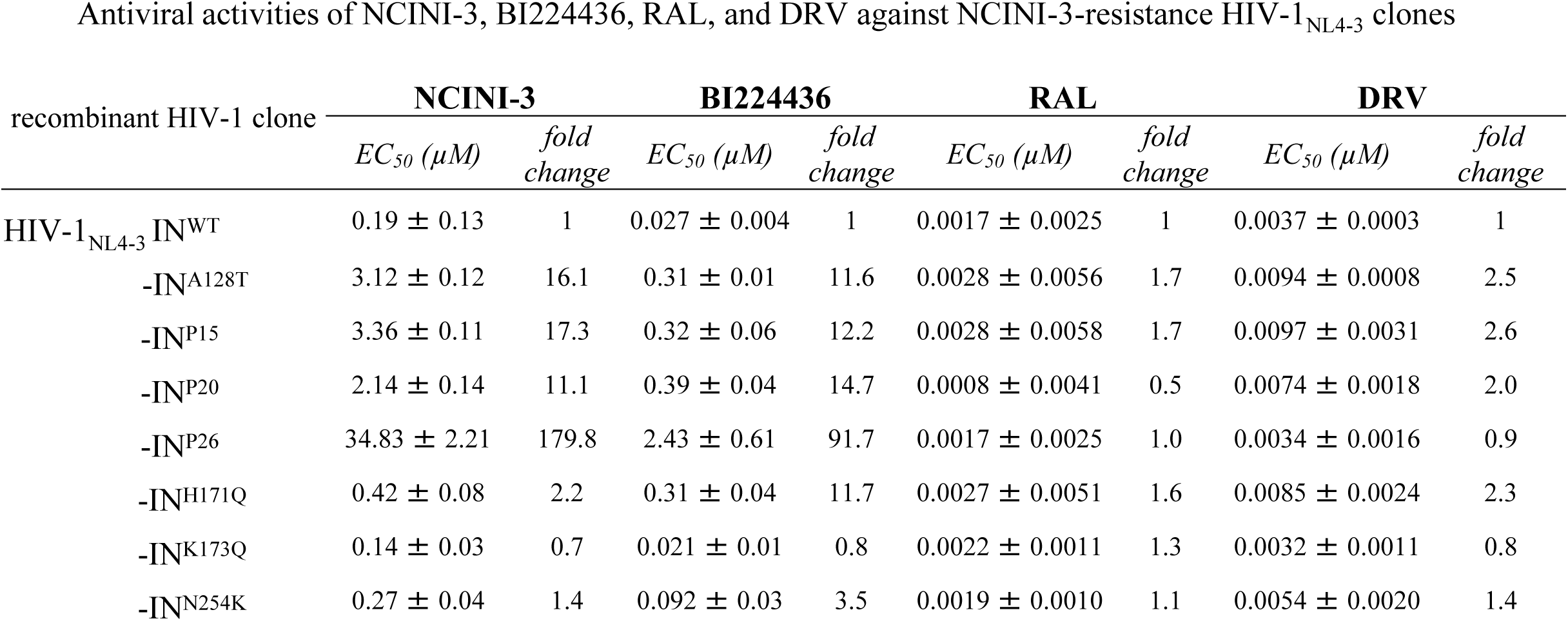
MT-4 cells (1 × 10^4^/ml) are exposed to 100 TCID_50_s of each infectious HIV-1 clone, and the inhibition of p24 production by the drug was used as the endpoint on day 7 in culture. The fold changes represent EC_50_ ratios against each HIV-1 clone to HIV-1 wild type. All assays are performed in duplicate, and the data shown are mean values (±SD) derived from the results of three independent experiments.

### Characteristics of recombinant IN proteins carrying NCINI-related resistance mutations

To examine how NCINI-3 resistance mutations acquire resistance against NCINI-3, we purified recombinant His-tagged IN proteins carrying mutations, such as A128T, H171Q, K173Q, N254K, P15, P20 and P26, respectively (IN^A128T^, IN^H171Q^, IN^K173Q^, IN^N254K^, IN^P15^, IN^P20^, and IN^P26^) expressed in *E. coli*, and analysed multimerization of the IN proteins. IN^WT^ multimerization appeared as two peaks of a tetramer and mixture of a dimer and monomer using size exclusion chromatography (SEC) (Figure 2A and Figure 2—figure supplement 2A, B). IN^H171Q^ and IN^N254K^ multimerization was very similar to that of IN^WT^, suggesting that H171Q and N254K mutations did not affect IN multimerization (Figure 2—figure supplement 2A, B), whereas IN^A128T^ multimerization was seen as a broader shoulder of the two peaks. IN^K173Q^ exhibited a moderately decreased tetramer peak. Further, IN^P15^, IN^P20^, and IN^P26^ proteins carrying the accumulated NCINI-3 resistance mutations completely shifted to one peak consisting of dimers and monomers. Next, we examined monomers, dimers, and tetramers of IN proteins cross-linking with BS3 using an immunoblotting assay (Figure 2B and Figure 2—figure supplement 2C). IN^WT^ and IN^P26^ were evaluated as a proportion of tetramers, dimers, and monomers. Multimer ratios of IN^P26^ (19.6 % dimers and 3.3 % tetramers) slightly decreased compared with those of IN^WT^ (22.9 % dimers and 6.5 % tetramers) (Figure 2B). In addition, results of the BiFC-IN system (***Nakamura et al., 2016***) carrying the NCINI-3 resistance mutations, previously reported to analyse IN multimerization in living cells, corroborated the SEC data (Figure 2C and Figure 2—figure supplement 2D, E, and Figure 2—table supplement 1).

**Figure 2.**
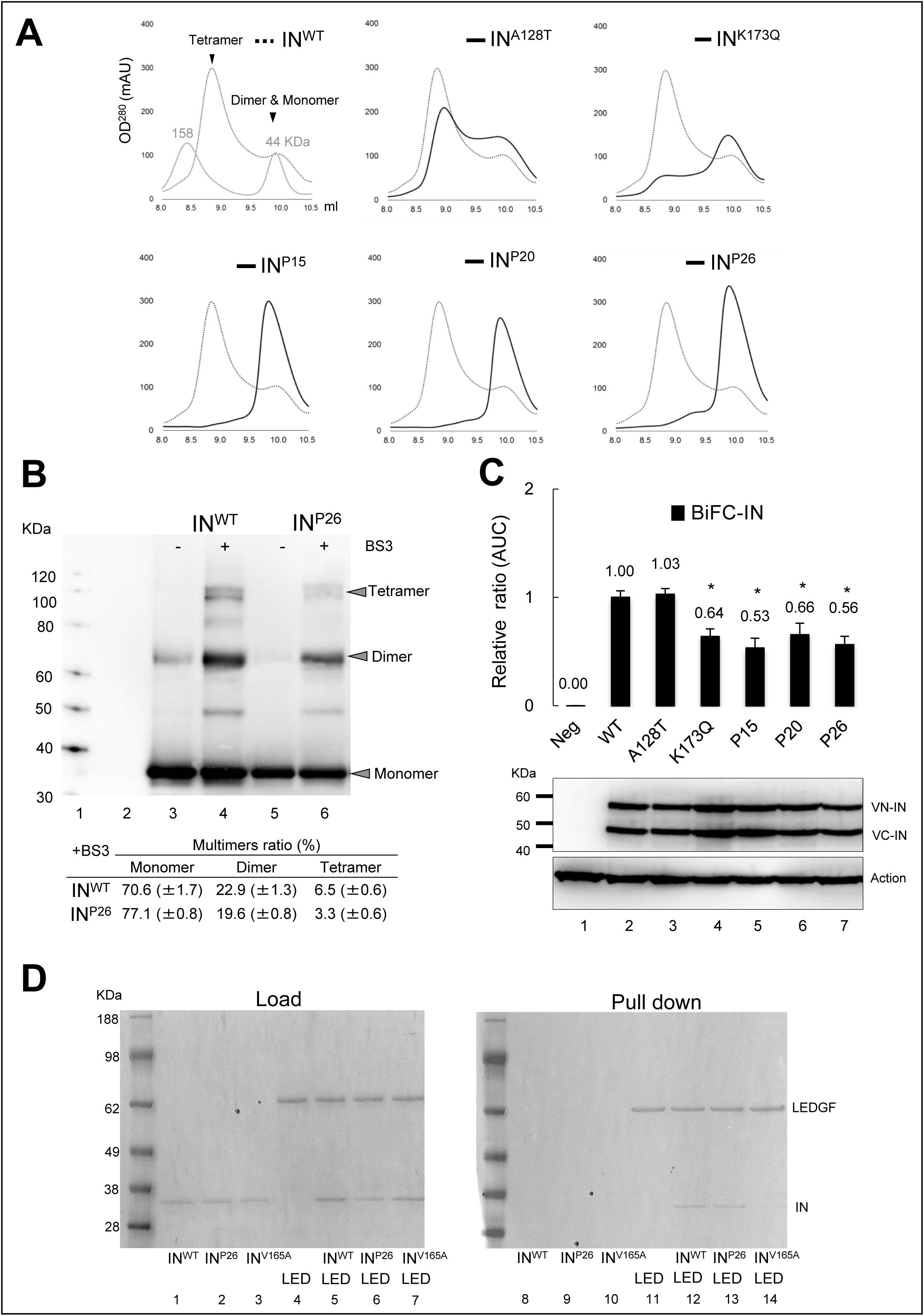
Characteristics of recombinant IN proteins carrying the NCINI-3 resistance mutations. (**A**) SEC analysis of IN proteins carrying the NCINI-3 resistance mutations. Elution profile of IN^WT^ proteins at 50 µM concentration is shown together with estimated IN multimers and two molecular weight standard proteins (158 and 44 KDa). Those of IN^A128T^, IN^K173Q^, IN^P15^, IN^P20^, and IN^P26^ proteins, respectively are shown compared with IN^WT^. (**B**) Multimers ratio of IN proteins (IN^WT^ and IN^P26^) cross-linking with BS3. Each multimer’s intensity of IN^WT^ and IN^P26^ measured by ImageJ is shown as multimers ratios (%). Molecular weight (MW) (Lane 1), buffer (Lane 2), IN^WT^ and IN^P26^ (Lanes 3 and 5), and IN^WT^ and IN^P26^ cross-linked with BS3 at 0.05 mM (Lanes 4 and 6) are shown by immunoblotting with anti-IN antibody. The ratios represent mean values (±standard deviation, SD) derived from immunoblotting results of three independent experiments. (**C**) Results of BiFC-IN system carrying the NCINI-3 resistance mutations. Fluorescence intensity derived from multimers of BiFC-IN^A128T^, -IN^K173Q^, -IN^P15^, -IN^P20^, and -IN^P26^ in HEK293T cells measured by FACS is shown as a ratio of area under the curve (AUC) compared to BiFC-IN^WT^. The AUC ratios represent mean values (±SD) derived from FACS results of three independent experiments. BiFC-IN system are summarized in ref. (***Nakamura T et al., 2016***) and Figure 2—figure supplement 2D, E, and Figure 2—table supplement 1. Statistical significance is examined using a parametric two tailed Student t-test, * P < 0.05. (**D**) LEDGF_His_ and F-IN pull down assay. Loading samples of F-IN^WT^, F-IN^P26^, F-IN^V165A^ (Lanes 1, 2, and 3), LEDGF_His_ (Lane 4), and LEDGF_His_ together with F-IN^WT^, F-IN^P26^, and F-IN^V165A^, respectively (Lanes 5, 6, and 7) are shown at left panel. Pull down samples of F-IN^WT^, F-IN^P26^, F-IN^V165A^ (Lanes 8, 9, and 10), LEDGF_His_ (Lane 11), and LEDGF_His_ incubated for 30 min with F-IN^WT^, F-IN^P26^, and F-IN^V165A^ (Lanes 5, 6, and 7), respectively after His-tag column selection are shown at right panel.

Next, to understand whether under-multimerized IN affects its interaction with LEDGF/p75, we performed a pull-down assay between His-tagged LEDGF/p75 and Flag-tagged IN proteins carrying P26 or V165A mutation reported as a Class II mutant which fails to interact with LEDGF/p75 (***Lu et al., 2004***). As shown in Figure 2D, IN^V165A^ as a negative control failed to interact with LEDGF/p75, whereas IN^P26^ proteins with IN under-multimerization were able to interact with LEDGF/p75 as well as IN^WT^. The accumulated NCINI-3 resistance mutations decreased IN multimerization. Specifically, P15, P20, and P26 mutations including at least A128T and K173Q mutations severely reduced IN multimerization, suggesting that IN under-multimerization may be one of the mechanisms used by HIV-1 to escape from the anti-HIV-1 effect of NCINIs.

### Characteristics of recombinant HIV-1NL4-3 clones carrying NCINI-related resistance mutations

Several potent NCINIs induce IN over-multimerization, leading to the production of immature viruses, but are not related to Gag and Gag-Pol proteolytic processing (***Jurado et al., 2013***; ***Sharma et al., 2014***) (Figure 3—figure supplement 3A).

In order to investigate characteristics of HIV-1 carrying the NCINI-3 resistance mutations with IN under-multimerization, we examined the process of viral production including Gag and Gag-Pol proteolytic processing, and viral maturation of the recombinant HIV-1 IN^K173Q^, -IN^P15^, -IN^P20^, and -IN^P26^ clones. As shown in Figure 2, K173Q, P15, P20, and P26 mutations significantly decreased IN multimerization; however, viral production levels (p24 values) of wild type HIV-1_NL4-3_ (HIV-1 IN^WT^) and such recombinant HIV-1 IN clones with IN under-multimerization were very similar at 72 h post-transfection (Figure 3—figure supplement 3B). Gag and Gag-Pol proteolytic processing of the recombinant HIV-1 IN clones with IN under-multimerization, analysed by immunoblotting in the presence of NCINI-3 or saquinavir (SQV, an HIV-1 protease inhibitor inhibits Gag-pol proteolytic processing), was similar to that of HIV-1 IN^WT^ (Figure 3A, Figure 3—figure supplement 3C), suggesting that the recombinant HIV-1 clones with IN under-multimerization had normal viral production, and Gag and Gag-Pol proteolytic processing.

**Figure 3.**
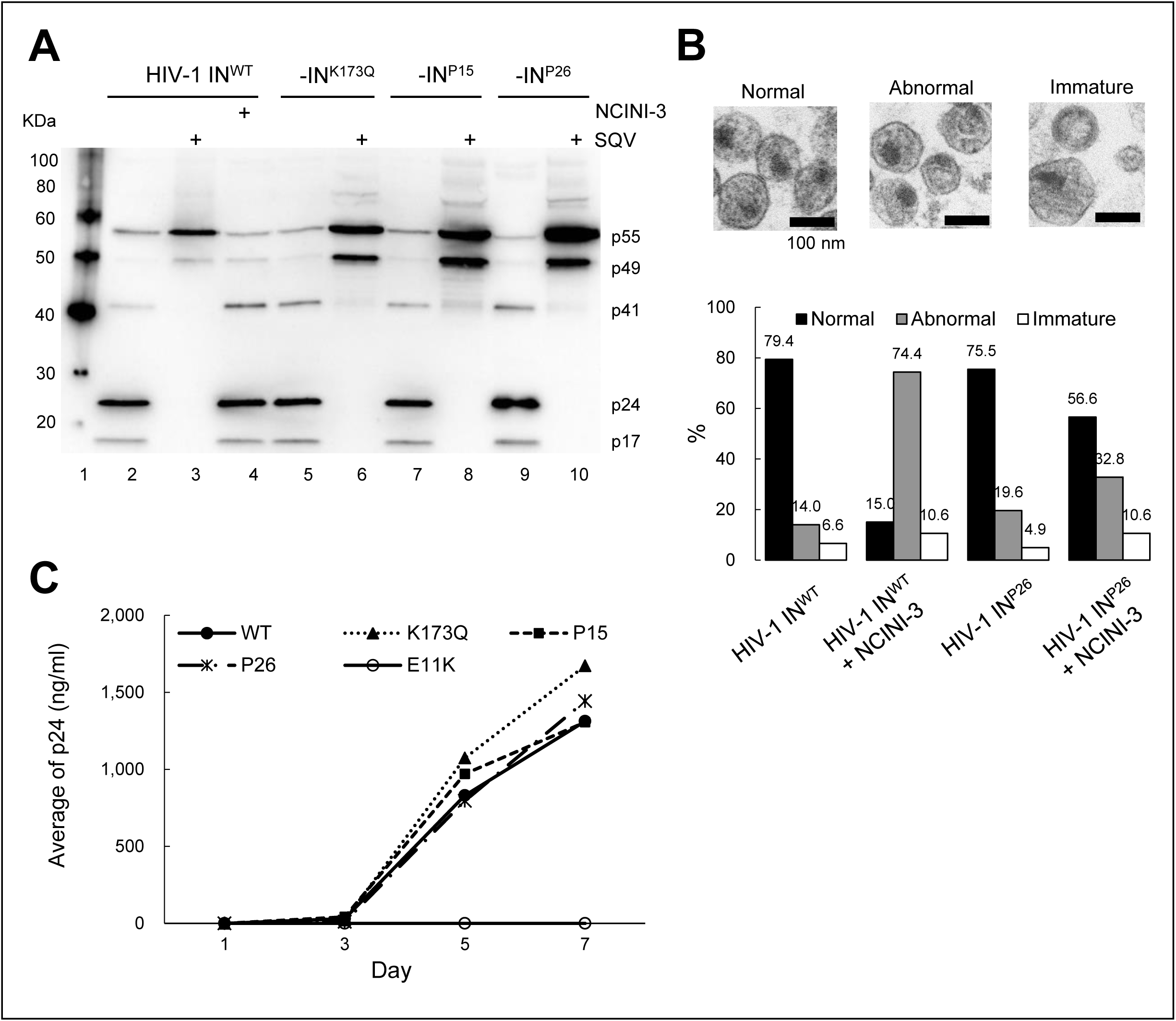
Characteristics of recombinant HIV-1 IN clones carrying the NCINI-3 resistance mutations with IN under-multimerization. (A) Gag proteolytic processing products of wild type and recombinant HIV-1_NL4-3_ IN^K173Q^, -IN^P15^, and -IN^P26^ clones, respectively. The HIV-1 IN clones produced by transfection of pNL_4-3_ encoding the NCINI-3 resistance mutations in the presence (Lanes 3, 6, 8, and 10) or absence (Lanes 2, 5, 7, and 9) of 2 µM SQV, or in the presence of 20 µM NCINI-3 (Lane 4). Gag proteolytic processing products normalized with p24 levels of the HIV-1 IN clones are visualized by immunoblotting with anti-HIV-1 p55 + p24 + p17 antibody. (**B**) Morphologies of HIV-1 IN^WT^ and HIV-1 IN^P26^ clone in the presence or absence of 20 µM NCINI-3 using TEM. Representative images of normal, abnormal, and immature HIV-1 particles (Magnification, 168,000 x scale bar, 100 nm) are shown at upper panel. Over 100 numbers of the HIV-1 particles are examined from several images of each HIV-1 sample, and percentages of HIV-1 morphology classified as normal, abnormal, and immature are shown in lower graph. (**C**) Replication kinetics of HIV-1 -IN^WT^ -IN^K173Q^, -IN^P15^, -IN^P26^, and -IN^E11K^ clones, respectively. MT-4 cells are exposed to the HIV-1 preparation normalized with the p24 levels, and production of the HIV-1 clones from the MT-4 cells is monitored at day 1, 3, 5, and 7 by p24 ELISA.

Besides, morphologies of HIV-1 IN^WT^ and HIV-1 IN^P26^ clone in the presence or absence of NCINI-3 were observed using a transmission electron microscope (TEM). HIV-1 particles were classified by viral structures under three phenotypes, normal mature conical core, abnormal (or empty) core, and immature (Figure 3B) as previously reported (***Jurado et al., 2013***; ***Desimmie et al., 2013***; ***Fontana et al., 2015***). The morphology of HIV-1 IN^WT^ included 79.4 % normal, 14.0 % abnormal, and 6.6 % immature structures. By addition of NCINI-3, abnormal morphology of HIV-1 IN^WT^ was identified in 74.4 % which indicated RNP translocation from the capsid core. On the other hand, the morphology of HIV-1 IN^P26^ clone with IN under-multimerization was similar to that of HIV-1 IN^WT^ and exhibited resistance against NCINI-3 (56.6 % normal, 32.8 % abnormal, and 10.6 % immature structures). Finally, we examined replication fitness of HIV-1 IN^WT^, recombinant HIV-1 IN clones with IN under-multimerization, and HIV-1 IN^E11K^ clone carrying E11K mutation with reported to impair IN tetramerization (***Hare et al., 2009***). The HIV-1 IN clones with IN under-multimerization infected in MT-4 cells showed robust replication fitness similar to that of HIV-1 IN^WT^ up to day 7 except HIV-1 IN^E11K^ clone (Figure 3C and Figure 3—table supplement 1). Thus, IN under-multimerization induced by NCINI-3 resistance mutations in this study did not significantly affect viral production including Gag and Gag-Pol processing, and viral maturation in the late stage of HIV-1 life cycle.

### Effect of certain mutations on stability and folding of IN catalytic core domain

It has been reported that NCINIs bind to two pockets formed in CCD proteins (***Christ et al., 2010***), carrying a solubilizing AA substitution, F185K (CCD^F185K^) (***Jenkins et al., 1995***). To examine whether NCINI-3 resistance mutations affect the structure of CCD, we produced and purified CCD^F185K^ proteins carrying NCINI-3 resistance mutations (CCD^A128T^, CCD^H171Q^, CCD^K173Q^, and CCD^P15^) (Figure 4—figure supplement 4A). It appeared that peak CCD^WT^ eluted at roughly the same monomer size as CCD—17 kDa (Figure 4A) by SEC in our method condition. CCD^H171Q^ which did not reduce the full-length IN multimerization (Figure 2—figure supplement 2B) eluted at the same size fraction as CCD^WT^ (Figure 4A). Notably, CCD^A128T^ and CCD^K173Q^ proteins, which reduced full-length IN multimerization, eluted at a decreased size compared to CCD^WT^, and CCD^P15^ eluted at a further decreased size, suggesting that A128T, K173Q, and P15 mutations may affect protein folding of the CCD monomer (Figure 4A).

**Figure 4.**
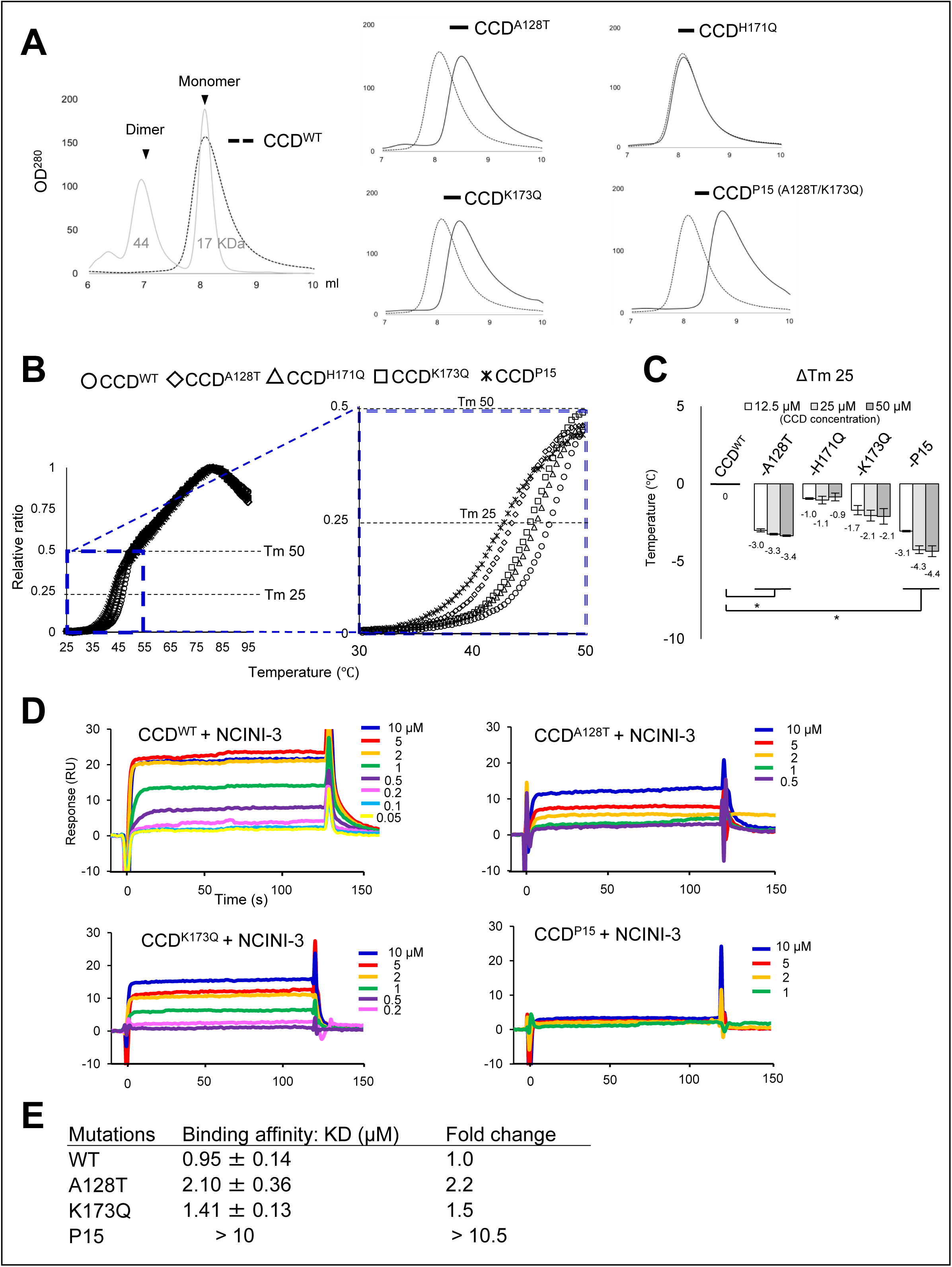
Effect of the NCINI-3 resistance mutations on stability and folding of CCD. (**A**) SEC analysis of the CCD proteins carrying the NCINI-3 resistance mutations together with the solubilizing F185K substitution. Elution profile of CCD^WT^ carrying is shown with two molecular weight standard (44 and 17 KDa) and estimating size position of CCD dimers and monomers, respectively. Those of CCD^A128T^, CCD^K173Q^, and CCD^P15^, respectively at 50 µM concentration are shown compared with CCD^WT^. (**B**) DSF analysis of CCD proteins carrying the NCINI-3 resistance mutations. Fluorescence of CCD^WT^ (○), CCD^A128T^ (◊), CCD^H171Q^ (△), CCD^K173Q^ (□), and CCD^P15^ (*) is detected and plotted as relative ratio at the temperature range from 25 to 95°C at left panel. Expanded plots at the low temperature range from 30 to 50°C are shown at right panel. (**C**) ΔTm 25 indicates Tm 25 differences between CCD^WT^ and each CCD^A128T^, CCD^H171Q^, CCD^K173Q^, and CCD^P15^ at 12.5, 25, and 50 µM CCD concentration, respectively. Statistical significance of ΔTm 25 was examined using the two tailed Student t-test. * P < 0.05. (**D**) SPR analysis of NCINI-3 interactions with CCD^WT^, CCD^A128T^, CCD^K173Q^, and CCD^P15^, respectively. SPR kinetics for NCINI-3 at indicated concentrations binding to each CCD^WT^, CCD^A128T^, CCD^K173Q^, and CCD^P15^ are shown. Data represents derived from two independent experiments. (**E**) Binding affinities (KD values) of NCINI-3 to CCD^WT^, CCD^A128T^, CCD^K173Q^, and CCD^P15^, and fold changes indicate KD value ratios of each CCD proteins to CCD^WT^. The KD values represent mean values (±SD) derived from two independent experiments.

In addition, to examine thermal stability of CCD^A128T^, CCD^H171Q^, CCD^K173Q^, and CCD^P15^, we performed differential scanning fluorimetry (DSF) (***Niesen et al., 2007***). The melting temperature value for proteins generally represents the temperature at which the protein is 50% folded (Tm 50). As shown in Figure 4B and Figure 4—figure supplement 4B, Tm 50 of all CCD proteins did not indicate significant differences. On the other hand, we specially noticed Tm 25 value, indicating the temperature at which CCD proteins are 25% folded upon heating. Tm 25 of CCD^A128T^, CCD^K173Q^, and CCD^P15^ proteins, except CCD^H171Q^, slightly decreased compared with that of CCD^WT^, depending on the concentration of the CCD proteins (from 12.5 to 50 µM) (Figure 4C), suggesting that CCD^A128T^, CCD^K173Q^, and CCD^P15^ were unstable at lower temperature from 35 to 45°C compared with CCD^WT^. Moreover, the surface hydrophobic sites of CCD^A128T^, CCD^K173Q^, and CCD^P15^ at 37°C slightly increased as shown in Figure 4—figure supplement 4C, D. These results suggest that A128T, K173Q, and P15 mutations affect thermal stability and folding of CCD proteins in the region of body temperature, possibly changing the conformation of CCD^A128T^, CCD^K173Q^, and CCD^P15^, respectively. Furthermore, to investigate the binding affinity of NCINI-3 to each CCD^A128T^, CCD^K173Q^, and CCD^P15^ proteins, we used Surface Plasmon Resonance (SPR). Binding affinity (KD value) of NCINI-3 to CCD^WT^ was 0.95 µM, while that of CCD^A128T^ and CCD^K173Q^ decreased by 2.10 µM (2.2-folds) and 1.41 µM (1.5-folds), respectively (Figure 4D, E). Further, CCD^P15^ carrying the double mutations (A128T/K173Q) did not interact under 10 µM NCINI-3 (Figure 4D, E).

These results indicate that such conformational alterations of CCD^A128T^, CCD^K173Q^, and CCD^P15^ proteins may also affect NCINI binding indirectly.

### The P15 mutation changes the monomer-monomer interface of the CCD dimer

It has been reported that NCINI-related compounds bind to CCD proteins containing F185K substitution by crystallographic analysis (***Dyda et al., 1994***; ***Christ et al., 2010***; ***Rouzic et al., 2013***). To confirm the conformational alteration of CCD^P15^, we analysed a crystal structure of CCD^P15^ using X-ray crystallography.

As shown in Figure 5A, the NCINI-related compound binds to the binding pocket in CCD^WT^ (PDB: 4GW6) (***Feng et al., 2013***), while a phenolic side chain (Y99 position) in CCD^P15^ (PDB: 6L0C) shifts closer to the NCINI-binding pocket compared with that in CCD^WT^ (Figure 5B), potentially interfering with NCINI to CCD^P15^ binding. Additionally, A128T mutation also affects the positioning of the NCINI as previously reported (***Feng et al., 2013***) (Figure 5—figure supplement 5A). Interestingly, the phenolic side chain of Y99 residue in a CCD^WT^ monomer (CCD1^WT^) participates in intermolecular interactions with two hydrogen bonds (H-bonds) to side chains of E87 and Q177, respectively, in the other CCD^WT^ monomer (CCD2^WT^) (Figure 5A). However, the interaction with Y99 in CCD1^P15^ reduces one H-bond with the side chain of H171 in CCD2^P15^; specifically, K173Q forms new intramolecular interactions with V88 and Q177 residues in CCD2^P15^ (Figure 5B, C). Such changes at the monomer-monomer interface in CCD^P15^ cause an unstable CCD^P15^ dimer compared with CCD^WT^, which may affect CCD^P15^ conformation including NCINI pocket binding.

**Figure 5.**
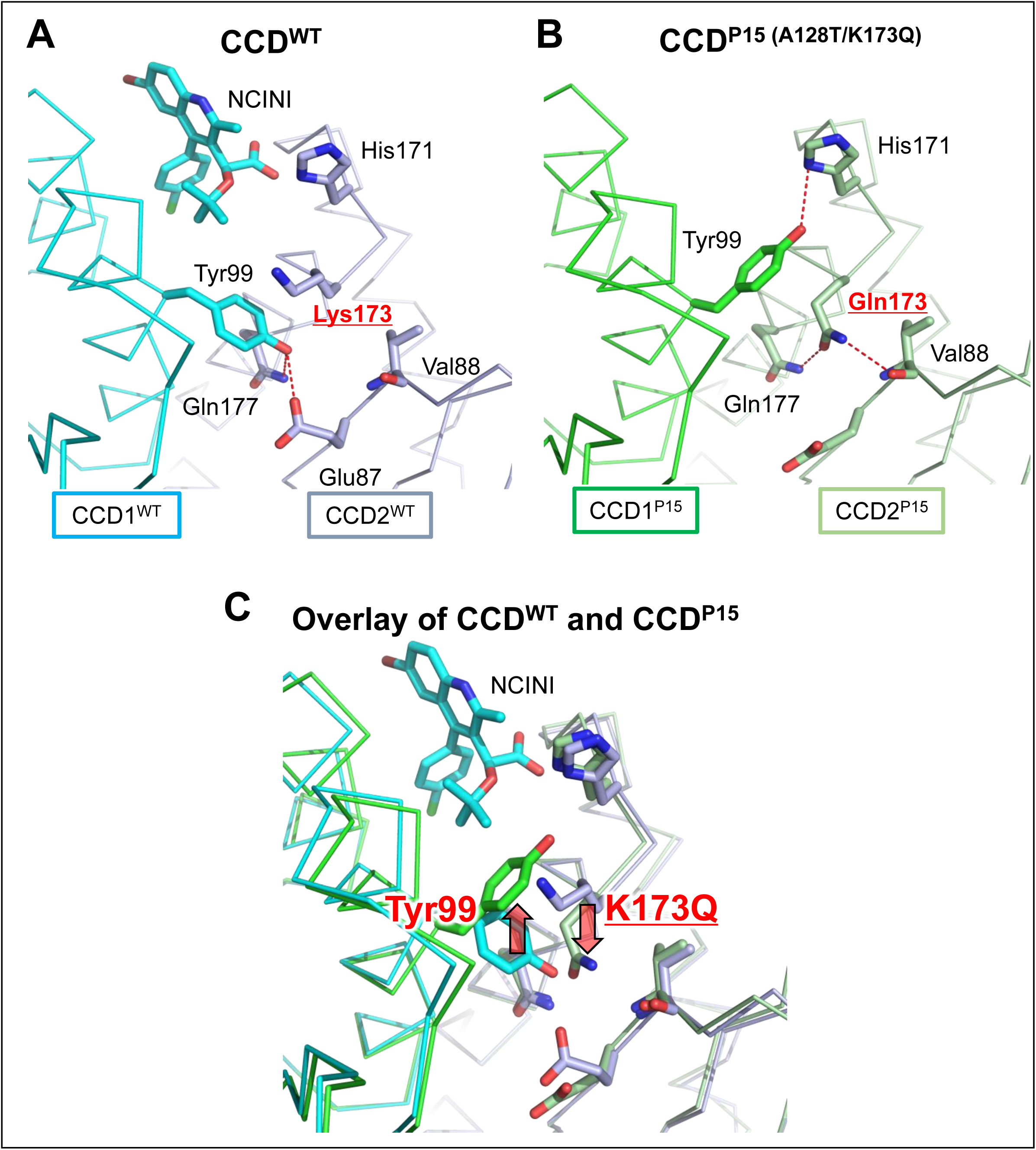
The P15 mutation changes the monomer-monomer interface of the CCD dimer. (**A**) The X-ray crystal structure of CCD^WT^ dimer (PDB: 4GW6). NCINI-related compound interacts with the binding pocket formed from CCD1^WT^ and CCD2^WT^ monomers colored in cyan and grey, respectively. Intermolecular hydrogen bonds between CCD1^WT^ and CCD2^WT^ are shown by red dashed lines. Locations of side chains for Y99 and K173 residues in the monomer-monomer interface of CCD^WT^ are indicated. (**B**) The X-ray crystal structure of CCD^P15^ dimer (PDB: 6L0C). CCD1^P15^ and CCD2^P15^ monomers are colored in green and light green, respectively. Intermolecular and intramolecular hydrogen bonds in CCD^P15^ dimers are shown by red dashed lines. Locations of side chains for Y99 and Q173 residues in the CCD^P15^ are indicated. (**C**) Overlay of the crystal structures of CCD^WT^ and CCD^P15^ binding to the NCINI-related compound.

These results suggest that the P15 mutation (A128T/K173Q) directly inhibits interaction of NCINIs with IN by shifting the bulky side chain of Y99, and may indirectly disrupt NCINIs binding by altering CCD^P15^ conformation, resulting in reduced full-length IN multimerization (IN under-multimerization) as a unique mechanism of HIV-1 escaping NCINI activity.

### HIV-1 RNA can stabilize and restore IN^P26^ multimerization

Over-multimerization of IN, directed by NCINIs, fails to interact with HIV-1 RNA, leading to production of immature, non-infective HIV-1 particles (Figure 3B) (***Kessl et al., 2016***).

To examine whether IN under-multimerization affects direct interaction with HIV-1 RNA, we analysed the direct interaction between purified Hisx6-tagged IN^P26^ proteins and biotinylated transactivation response (TAR) element, 54 RNA nucleotides, using alpha screen assay 1 (***Demeulemeester et al., 2012***; ***Kessl et al., 2016***).

In this assay, we employed recombinant IN^RAKA^ proteins carrying R269A/K273A mutations, with reported normal IN multimerization but loss of interaction with HIV-1 RNA, as a negative control (***Kessl et al., 2016***) (Figure 6—figure supplement 6A, B). FRET signals of each IN^WT^ and IN^P26^ at 10 nM were sufficiently increased by the addition of TAR-RNA concentration-dependent up to 10 nM (Figure 6A). Interestingly, the average FRET signals of IN^RAKA^ were seen at a low level in only 10 nM of TAR-RNA, whereas those of IN^P26^ were detected at a higher level than those of IN^WT^ at 10 nM (Figure 6A), suggesting that the binding ability of IN^P26^ to HIV-1 RNA was maintained or potentially reinforced.

**Figure 6.**
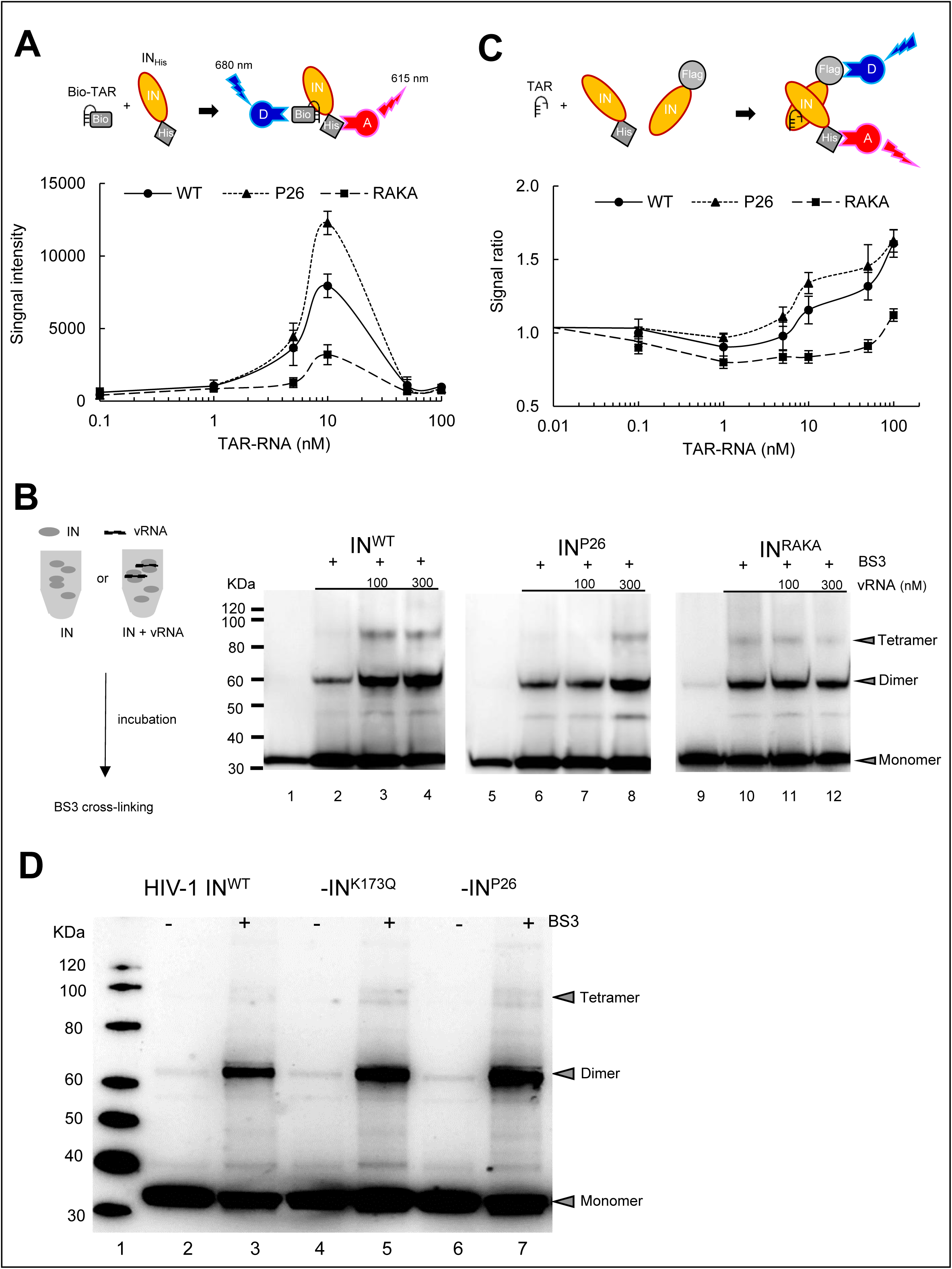
HIV-1 RNA stabilizes and restores IN under-multimerization. (**A**) AlphaScreen assay 1 for direct binding between Bio-TAR and IN_His_. Schematic illustration indicates that streptavidin coated Donor (D) and anti-His Accepter (A) beads bind to Bio-TAR and IN_His_ complex at upper panel. FRET signal intensity between IN proteins (IN^WT^, IN^P26^, and IN^RAKA^ at 10 nM) and Bio-TAR at various concentration is shown at left panel. Data represents mean values (±SD) derived from results of three independent experiments. (**B**) Multimerization of purified His-tagged IN^WT^, IN^P26^, and IN^RAKA^ proteins cross-linking with BS3. Schematic illustration of the method is shown at left panel. IN^WT^, IN^P26^, and IN^RAKA^ proteins at 300 nM are incubated with 100 and 300 nM of HIV-1 RNA (1-850 bp), respectively. IN^WT^, IN^P26^, and IN^RAKA^ proteins cross-linked with 0.05 mM of BS3 (Lanes 2, 3, 4, 6, 7, 8, 10, 11, and 12) are employed 7.5 µl, and without BS3 (Lanes 1, 5, and 9) employed 3 µl, and analysed by immunoblotting with anti-HIV-1 IN antibody. (**C**) AlphaScreen assay 2 for indirect IN multimerization between F-IN and IN_His_. Schematic illustration indicates that anti-Flag coated D and anti-Nickel coated A beads bind to a dimer of F-IN and IN_His_. FRET signal intensity between IN_His_ and F-IN proteins (IN^WT^, IN^P26^, and IN^RAKA^, respectively) with various concentration of TAR-RNA is shown as a signal ratio to the signal without TAR-RNA. Data represents mean values (±SD) derived from three independent experiments. (**D**) Profile of IN multimerization in HIV-1_NL4-3_ IN^WT^, -IN^K173Q^, and -IN^P26^ viral particles. HIV-1 IN^WT^, -IN^K173Q^, and -IN^P26^ clones are purified by ultracentrifugation. MW (Lane 1), the high concentrated HIV-1 samples stripped the viral membrane are cross-linked with (Lanes 3, 5, and 7) or without (Lanes 2, 4, and 6) BS3, and visualized by SDS-PAGE with anti-HIV-1 IN antibody.

Next, in order to confirm the effect of HIV-1 RNA on IN multimerization, we investigated IN cross-linking with BS3 in the presence of HIV-1 transcriptional RNA, 850 RNA nucleotides, using an immunoblotting assay. As shown in Figure 6B, tetramers of IN^WT^ at 300 nM were observed in the presence of 100 nM of HIV-1 RNA, while those of IN^P26^ were observed only in the presence of 300 nM of HIV-1 RNA (IN : RNA = 1 : 1). However, tetramers of IN^RAKA^ were not changed despite increasing of HIV-1 RNA. Moreover, to quantitatively examine IN multimerization in the presence or absence of TAR-RNA, we generated alpha screening assay 2 with Flag- and Hisx6-tagged IN proteins. FRET signal ratios in 10 nM of IN^WT^ and IN^P26^, respectively, increased by concentration-dependent TAR-RNA (from 0.01 to 100 nM) except IN^RAKA^ (Figure 6C). These data suggested that IN multimerization is stabilized and increased in the presence of HIV-1 RNA.

Finally, we investigated IN^WT^, IN^K173Q^, and IN^P26^ multimerization of the recombinant HIV-1 clones using cross-linking with BS3 immunoblotting assay after membrane stripping with 0.5% Triton X in PBS. Multimer ratios of IN^K173Q^ and IN^P26^ in the viral particles were comparable to those of IN^WT^ (Figure 6D and Figure 6—figure supplement 6C), indicating that IN under-multimerization induced by the mutations was relieved in the viral particle containing HIV-1 RNA.

Taken together, IN proteins with IN under-multimerization via NCINI-resistant mutations can bind to HIV-1 RNA and recover the IN multimerization during HIV-1 maturation in the viral particle. This mechanism allows NCINI-resistant HIV-1 to escape from NCINI action and propagate within the cells (Figure 7).

**Figure 7.**
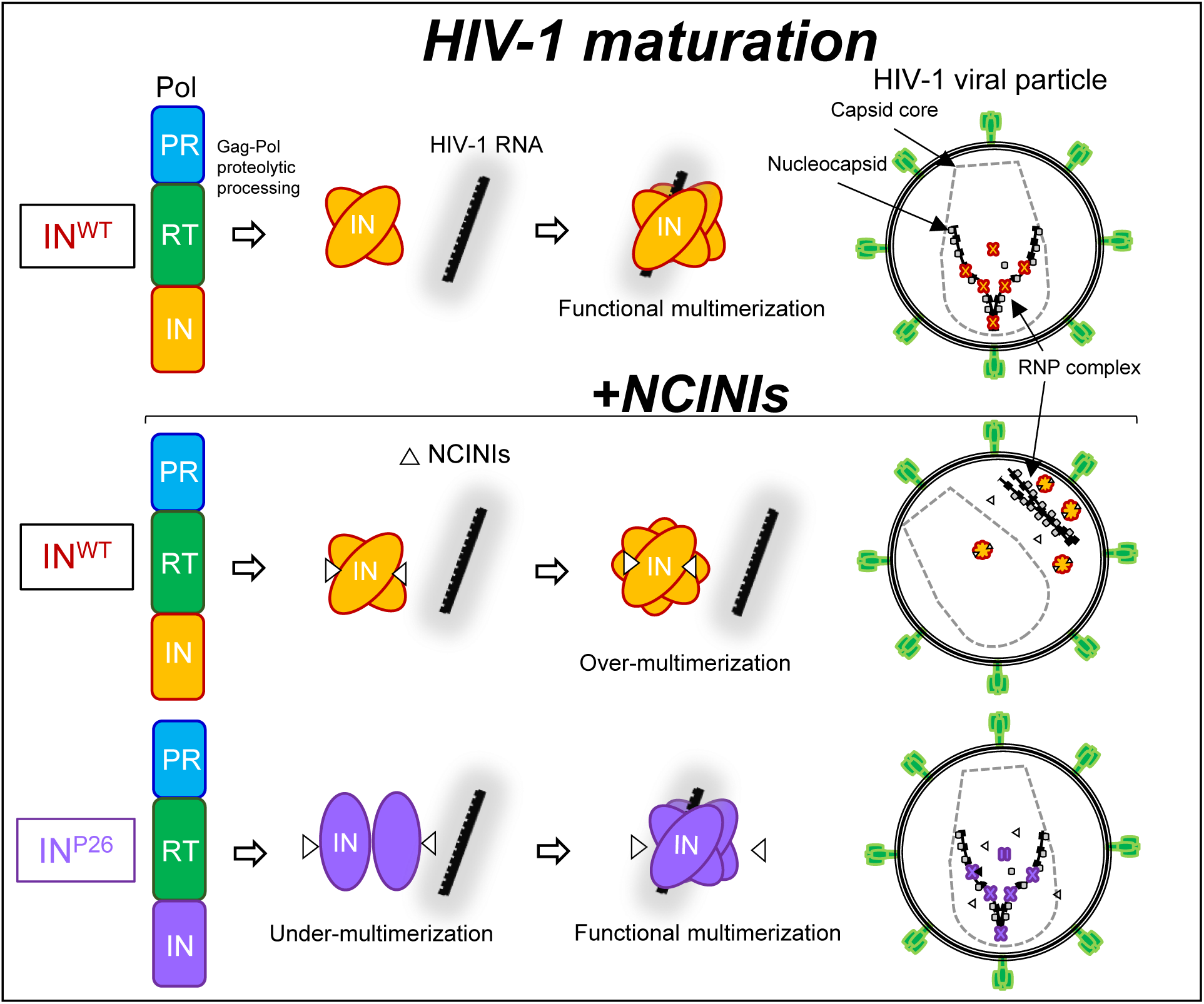
Graphical article during HIV-1 maturation. Schematic illustration focuses on the interaction of IN multimerization with HIV-1 RNA, and the location of the RNP in the viral particles during HIV-1 maturation from Gag-Pol proteolytic processing. Upper illustration indicates that normal interaction of functional IN multimerization with HIV-1 RNA results in mature HIV-1. Middle illustration shows that over-multimerized IN proteins induced by NCINIs are not able to interact with HIV-1 RNA, producing immature HIV-1 in which the RNP complex translocated from the capsid core. Lower illustration indicates that under-multimerized IN^P26^ can escape from NCINIs binding and is restored to IN^P26^ with functional multimerization by HIV-1 RNA binding, producing mature HIV-1 in the presence of potent NCINIs.

## Discussion

In the present study, we found that IN undergoes a novel conformational alteration to escape NCINI activity. HIV-1 easily acquired drug-resistant mutations against NCINIs in our selection experiments (Figure 1). A128T was reported as a typical and critical NCINI-related resistance mutation (***Feng et al., 2013***). We also observed that A128T emerged early at P15 passage together with K173Q in the presence of NCINI-3, and was acquired in all NCINI-related resistant HIV-1 variants when selected in the presence of NCINI-1, -2, or -3 (Figure 1). Notably, in our HPLC data, IN^A128T^ multimerization slightly changed, and K173Q mutation moderately reduced IN multimerization. Further, P15, P20, and P26 containing A128T/K173Q mutations severely reduced IN tetramers to monomer and dimer subunits. This implicates acquisition of a combination of A128T and K173Q mutations for the reduction of IN multimerization (IN under-multimerization) as an escape mechanism of HIV-1.

It was reported that CCD dimers formed at the central core are material component of full-length IN tetramers with HIV-1 nucleotides (***Hare S et al., 2010; Passos et al., 2017***). However, without HIV-1 nucleotides, the P15 mutation changed the folding and stability of CCD monomers, possibly inducing unstable CCD dimerization, altering CCD conformation which forms NCINI binding sites. Such conformational changes could induce severe reductions of full-length IN multimerization before IN binding to HIV-1 RNA. IN^P26^ multimerization was recovered in the presence of HIV-1 RNA (Figure 6), and HIV-1 IN^P26^ clone showed replication fitness similar to HIV-1 wild type (Figure 3C and Figure 3—table supplement 1), suggesting that IN proteins carrying A128T/K173Q mutations with IN under-multimerization still retained intrinsic IN function.

Based on the EC_50_ ratios of these resistance mutations (Table 1 and Table 1—table supplemental 1), HIV-1 IN^A128T^ exhibited strong resistance to NCINI-3 and BI224436, suggesting that A128T represents a critical mutation against NCINI action in line with previously reported data (***Feng et al., 2013***). On the other hand, HIV-1 IN^K173Q^ clone carrying K173Q, a unique mutation which moderately reduces IN multimerization, showed resistance (over 10 µM) against weak NCINIs (NCINI-1 and NCINI-2) (Table 1—table supplemental 1), but no observed resistance against potent NCINIs (NCINI-3 and BI224436). By the addition of A128T, HIV-1 IN^P15^ clones acquired severe resistance to the potent NCINIs. Collectively, K173Q mutation moderately decreased IN multimerization and reduced direct binding of NCINI-3 (Figure 2 and 3), but, EC_50_ of HIV-1 IN^K173Q^ clone did not indicate critical resistance compared with that of HIV-1 IN^A128T^ clone, suggesting that K173Q plays a secondary role among the NCINI-related resistance mutations. The H171Q mutation did not affect IN multimerization (Figure 2—figure supplement 2B), but showed resistance against NCINIs (Table 1). H171T mutation at the same AA position was reported as a NCINI-related resistance mutation (***Slaughter et al., 2014***). According to the report, H171T did not affect IN multimerization and showed resistance to NCINI-related compounds. The characteristics of H171T were similar to those of H171Q, suggesting that HIV-1 IN^H171Q^ and IN^H171T^ clones possibly express different phenotypes compared to the HIV-1 IN^K173Q^ clone. T124N/T174I resistance mutations against a pyridine-based inhibitor, KF116, was also reported to disrupt IN multimerization (***Hoyte et al., 2017***). The T124N/T174I mutations decreased Gag-Pol proteolytic processing, viral maturation, and viral infectivity of the HIV-1 IN^T124N/T174I^ variant compared with those in wild type HIV-1. It seems that the T124N/T174I mutation, which induced abnormal IN multimerization, confers a phenotype similar to that of class II IN mutant HIV-1 (***Engelman et al., 1999***); however, A128T/K173Q showed phenotypes clearly different from that of the class II IN mutant.

We found a unique escape reaction of HIV-1 from NCINIs using mutation analysis, although NCINIs are not yet approved by the FDA. NCINI-related resistance mutations in vivo are unknown, which present the major limitations of our study. With regards to the interaction of IN proteins with HIV-1 RNA, it is likely that the interaction of HIV-1 RNA with under-multimerized IN^P26^ may be stronger than that with IN^WT^, however it is still unclear how IN^P26^ increased the interaction of HIV-1 RNA. HIV-1 nucleocapsid (NC) binds to HIV-1 RNA, constituting RNP, while IN also binds to HIV-1 RNA at different positions from NC to keep RNP in the capsid conical core (***Fontana et al., 2015***; ***Kessl et al., 2016***). It has yet to be shown how IN bindings to HIV-1 RNA maintains RNP in the capsid core during HIV-1 maturation.

In conclusion, investigation into the drug-resistant mutations associated with HIV-1 protein multimerization may facilitate elucidation of the molecular mechanisms of the protein functional multimerization, aiding in developing more potent anti-HIV-1 drugs and unique treatment strategies.

## Materials and Methods

### Cells

293T (JCRB cell bank, Japan) and HEK293T (ATCC, CRL-11268) cells were maintained in Dulbecco’s Modified Eagle’s Medium (DMEM; WAKO, Japan) supplemented with 10% fetal bovine serum (FBS; SIGMA-ALDRICH), penicillin (P+) (100 I.U/mL), and kanamycin (K+) (100 mg/mL) (MEIJI, Japan) at 37°C and 5% CO_2_. MT-2 and MT-4 cells (JCRB) were in RPMI-1640 medium (WAKO, Japan) supplemented with 10% FBS, P+, and K+ at 37°C and 5% CO_2_.

### Plasmids constructs

Full-length IN and CCD sequences derived from pNL_4-3_ were introduced to pET30a vectors (Novagen) with Hisx6 tagged at the C-terminus or N-terminus, respectively, producing pET30a IN-6xHis (IN_His_) or pET30a 6xHis-CCD (CCD_His_) by using In-Fusion® HD Cloning Kit (TAKARA Bio Inc., Japan). Flag-tagged IN (F-IN) was also introduced to pET30a vector at the N-terminus by the In-Fusion method, producing pET30a Flag-IN. LEDGF/p75 sequence derived from 293T cells was introduced to pET30a vectors (Novagen) with Hisx6 tagged at the N-terminus, producing pET30a His-LEDGF/p75 (LEDGF/p75_His_). The site-directed mutagenesis was performed using PrimeSTAR**^®^** Max (TAKARA) to introduce the NCINI-3 resistance single mutations (A128T, H171Q, K173Q, and N254K), E11K, V165A, and R269A/K273A into the pET30a IN_His_, producing pET30a IN^A128T^, IN^H171Q^, IN^K173Q^, IN^N254K^, IN^E11K^, IN^V165A^, and IN^RAKA^ vectors, respectively. The pET30a CCD_His_ containing a solubilizing F185K substitution was introduced the NCINI-3 resistance mutations, producing pET30a CCD^A128T^, CCD^H171Q^, and CCD^K173Q^ vectors. The accumulated NCINI-3 mutations of P15 (A128T/K173Q), P20 (A128T/K173Q/N254K), and P26 (A128T/H171Q/K173Q/N254K) were introduced to pET30a IN_His_ or CCD_His_ vector by In-Fusion method, producing pET30a IN^P15^, IN^P20^, IN^P26^, and pET30a CCD^P15^, respectively. The NCINI-3 resistance mutations were also introduced to pNL_4-3_ and pBiFC-IN vectors (***Nakamura et al., 2016***) by above the method.

### Proteins expression and purification

IN and CCD proteins were produced from the pET30a IN_His_ and CCD_His,_ respectively in E. coli Rosetta^TM^ (DE3) pLysS Competent Cells (Novagen) grown in LB medium supplemented with K+ and chloramphenicol (Cam+) at 37°C and induced with 1.0 mM Isopropyl β-D-1-thiogalactopyranoside (IPTG) for 4 h at 30°C. The bacterial cells were harvested and stored at −80°C. The pellets of IN_His_ were resuspended and sonicated in lysis buffer (50 mM HEPES-NaOH pH 7.4, 100 mM NaCl). The lysates were cleared by centrifugation for 15 min at 3,500 rpm at 4°C. The supernatants were removed, and the pellets of IN were resuspended in IN reservoir buffer (50 mM HEPES pH 7.4, 1 M NaCl, 7.5 mM CHAPS, 10 mM MgSO_4_, 10% glycerol) supplemented with 5 mM 2-mercaptoethanol (BME), 10 mM imidazole and 0.5 mM Phenylmethylsulfonyl fluoride (PMSF). The pellets of CCD were resuspended and sonicated in CCD reservoir buffer (50 mM HEPES pH 7.4, 500 mM NaCl, 7.5 mM CHAPS, 10 mM MgSO_4_, 5% glycerol) supplemented with 5 mM BME, 10 mM imidazole, and 0.5 mM PMSF. The lysates were precleared for 30 min at 15,000 rpm at 4°C, and filtered through a 0.45 μm filter. IN and CCD proteins were loaded onto a His-TALON column (TAKARA), and the columns were washed with IN wash buffer (20 mM HEPES pH 7.4, 1 M NaCl, 3.75 mM CHAPS) or CCD wash buffer (20 mM HEPES pH 7.4, 500mM NaCl, 3.75 mM CHAPS). IN or CCD proteins were eluted with IN elution buffer (50 mM HEPES ph7.4, 1 M NaCl, 7.5 mM CHAPS, 10 mM MgSO_4_, 10% glycerol, 300 mM imidazole) or CCD elution buffer (50 mM HEPES pH 7.4, 500 mM NaCl, 7.5 mM CHAPS, 10 mM MgSO_4_, 5% glycerol, 300 mM imidazole) using AKTAprime plus (GE Healthcare). IN or CCD fractions were concentrated using Amicon Ultra-30K or -10K device (Merck Millipore) in the each reservoir buffer. LEDGF/p75 was produced in E. coli RosettaTM (DE3) pLysS competent cells grown in LB medium supplemented with 2% ethanol, K+ and Cam+ at 37°C and induced with 0.5 mM IPTG for 5 h at 30°C. The bacterial cells were harvested and sonicated in LEDGF reservoir buffer (50 mM Tris-HCl pH 7.4, 500 mM NaCl, 5 mM BME) supplemented with 10 mM imidazole and 0.5 mM PMSF. The lysates were precleared for 30 min at 15,000 rpm at 4°C, and filtered. LEDGF/p75 was loaded onto the His-TALON column and the column was washed with LEDGF wash buffer (20 mM Tris-HCl pH 7.4, 500 mM NaCl). LEDGF/p75 was eluted with LEDGF elution buffer (50 mM Tris-HCl pH 7.4, 500 mM NaCl, 5mM BME, 300 mM imidazole). LEDGF were concentrated using Amicon Ultra-50K device in LEDGF reservoir buffer. F-IN was expressed as above the method of IN_His_ and purified by previous reports (***Kessl et al., 2012***). The proteins concentration was determined using a BCA Protein Assay Reagent Kit (Thermo Fisher Scientific) and stored at −80°C.

### Recombinant HIV-1 clones

293T cells (1.5 × 10^5^/ml) were seeded on TC 6-well plate (Greiner Bio-one) and incubated for 24 h at 37°C. Cells were transfected with 5 μg of pNL_4-3_ encoding NCINI-3 resistance mutations (A128T, H171Q, K173Q, N254K, P15, P20, and P26), or E11K in the IN region using Attractene Transfection Reagent (QIAGEN). Cells were washed at 24 h later, and cell supernatants containing viruses were collected after 48 h incubation. The HIV-1 clones were measured by HIV-1 p24 ELISA (LUMIPULSE® G1200; FUJIREBIO Inc., Japan) and normalized to determine viral concentration, and stored at −80°C.

### Western blotting

293T cells were plated on TC 6-well plates (1.5 × 10^5^/ml) and incubated for 24 h at 37◦C in 5% CO_2_. Cells were co-transfected with BiFC-IN vectors carrying NCINI-3 resistance mutations using Attractene and incubated for 48 h. Subsequently, the cells were lysed in lysis buffer (20 mM Pipes pH 7.0, 400 mM NaCl, 10% (w/v) sucrose, 0.5 mM dithiothreitol (DTT), 0.5% NP-40, Halt Protease Inhibitor Cocktail (Thermo Fisher Scientific) (***Cherepanov et al., 2003***). After centrifugation for 30 min at 15,000 rpm at 4°C, the samples were titrated using BCA Protein Assay Kit and stored at −80°C. Wild type and recombinant HIV-1 clones were filtered and purified by ultracentrifugation (at 35,000 rpm for 30 min) in 15% sucrose PBS, and normalized by the p24 levels and stored in PBS supplemented with Halt Protease Inhibitor Cocktail at −80°C. The samples were prepared in NuPAGE® LDS and reducing sample buffer (Thermo Fisher Scientific), and separated with SDS-PAGE (4–12% Bis-Tris Gel; Thermo Fisher Scientific) and transferred a nitrocellulose membrane. The samples were detected with anti-HIV-1 integrase antibody (abcam; ab66645), anti-HIV-1 p55 + p24 + p17 antibody (abcam; ab63917), anti-HIV1 p24 [39/5.4A] (abcam; ab9071), and second mouse or rabbit antibody (MBL co., LTD.), and anti-beta actin antibody (HRP conjugated) (abcam), and then visualized using SuperSignal WestPico Chemiluminescent Substrates (Thermo Fisher Scientific).

### Cross-linking IN proteins with BS3

Purified IN proteins were diluted at 300 nM final concentration in buffer (20mM HEPES pH 7.4, 100 mM NaCl, 2 mM DTT, 5 mM MgCl_2_, and 15 µM ZnCl_2_). Samples were mixed with BS3 (Thermo Scientific Pierce) at 0.05 mM final concentration, and incubated for 20 min at RT, and the reactions were quenched by the addition of mixed LDS and reducing sample buffer. The cross-linked proteins with BS3 were visualized by immunoblotting. In the case of IN in viral particles (***Balakrishnan et al., 2013***), the high concentrated wild type and recombinant HIV-1 clones were titrated, and treated with 0.5% triton X in PBS, and cross-linked with 0.05 mM BS3. The samples employed at 100 ng (p24 level) per wells were visualized by immunoblotting with anti-IN antibody and SDS PAGE (4–12% Bis-Tris Gel).

### Generation of drug resistant HIV-1 variants

Selection experiments of resistant HIV-1 variants were performed as previously described (***Amano et al., 2007***). In brief, MT-4 cells (1.0 × 10^5^/ml) were exposed to HIV-1_NL4-3_ (100 TCID_50_) and cultured in the presence of NCINIs, each at an initial concentration of each EC_50_. Viral replication in MT-4 cells was monitored by p24 levels of the culture medium at intervals of one week. The selection procedure was continued until the compound concentration over 10 µM. Proviral DNA sequences of resistant HIV-1 variants were extracted from the infected into MT4 cells using NucleoSpin^®^ Tissue (TAKARA). DNA sequences of the IN regions were amplified with 1^st^ primers (5’-AAA TTT AAA TTA CCC ATA CAA AAG GAA ACA TGG GAA GC-3’ and 5’-GGT CTG CTA GGT CAG GGT CTA CTT GTG TGC TAT ATC-3’) and following 2^nd^ nested primers (5’-AAA AGG AAA AAG TCT ACC TGG CAT GGG TAC CAG CAC AC-3’ and 5’-AGT CCT TAG CTT TCC TTG AAA TAT ACA TAT GG-3’), and then analysed using IN sequence primers (5’-AAA AGT TAT CTT GGT AGC AGT TCA TGT AGC CAG TGG-3’ and 5’-TT TAG TTT GTA TGT CTG TTG CTA TTA TGT CTA CTA TTC-3’).

### Biomolecular fluorescence complementation (BiFC) analysis

BiFC-IN system was performed as previously described (***Nakamura et al., 2016***) (Figure 2—table supplement 1). This assay was modified as following. In brief, HEK293T cells were seeded (2 × 10^4^/well) on Collagen I 96-well microplates (BD BioCoat^TM^) and incubated for 24 h at 37◦C in 5% CO_2_. The cells were co-transfected with pBiFC-IN vectors (VN-IN and VC-IN) carrying the NCINI-3 resistance mutations using Attractene and incubated for 48 h at 37◦C. Then Venus fluorescence in the cells was measured using a FACS (FACSVerse flow cytometer, BD Bioscience). Area under the curve (AUC) of the histogram which consists of the cell count and fluorescent intensity emitting from BiFC-IN was measured using ImageJ (Figure 2—figure supplement 2D). Data was compared as AUC ratios to AUC of BiFC-IN^WT^.

### IN and LEDGF/p75 binding assay

Pull down assay was performed to confirm the interaction of F-IN with LEDGF/p75_His_ (***Kessl et al., 2016***; ***Hoyte et al., 2017***). Briefly, IN (2 μM) and LEDGF/p75 (2 μM) in binding buffer (50 mM HEPES pH 7.0, 300 mM NaCl, 2 mM MgCl_2_, 2 mM BME, 5% Glycerol, and 0.2% (v/v) Nonidet P-40) supplemented with 20 mM Imidazole were incubated for 30 min at RT, and loaded on His 60 Ni resin (TAKARA) after equilibrating. The reaction mixture was incubated for 30 min at RT, and then were repeated washing by wash buffer (50 mM HEPES pH 7.0, 1M NaCl, 5 mM BME, and 5% Glycerol). The samples were eluted by the binding buffer supplemented with 300 mM imidazole, and visualized by Coomassie Brilliant Blue staining.

### HPLC

Recombinant IN_His_ proteins were examined with a TSKgel-SW_XL_G3000 column (TOSOH cor., Japan) at 1.0 ml/min in running buffer (20 mM HEPES pH 7.4, 750 mM NaCl, 3.75 mM CHAPS) (***Hare et al., 2009***). IN proteins (50 μM) in sample buffer (50 mM HEPES pH 7.4, 1 M NaCl, 7.5 mM CHAPS, 10 mM MgSO_4_, 50 µM ZnCl_2_, 2 mM DTT, 10% Glycerol) were subjected to Agilent Technologies 1220 Infinity LC system (TOSOH) for SEC, and were detected by OD at 280 nm. Recombinant CCD_His_ proteins were examined with a COSMOSIL 5Diol-120-II column (nakarai tesque, Japan) at 1.0 ml/min in running buffer (20 mM HEPES pH 7.4, 500 mM NaCl, 3.75 mM CHAPS). CCD proteins (50 μM) in sample buffer (50 mM HEPES pH 7.4, 500 mM NaCl, 7.5 mM CHAPS, 10 mM MgSO_4_, 5 mM BME, 5% Glycerol) were subjected for SEC. The column was calibrated with standard proteins containing Bovine gamma-globulin (158 KDa), Chicken ovalbumin (44 KDa), and Horse myoglobin (17 KDa). All the procedures were controlled at 4°C as possible.

### Crystallization, X-ray data collection and structure determination

Crystallization procedure was referred to a previous report (***Rouzic et al., 2013***). Briefly, crystallization was performed by the hanging drop vapor diffusion method using EasyXtal 15-Well Tools (QIAGEN). CCD^P15^ proteins carrying A128T, K173Q, and F185K mutations were expressed and purified as describe above. The solution buffer was changed to 50 mM MES-NaOH pH 5.5, 50 mM NaCl, and 5 mM DTT using an Amicon Ultra-10K device (Millipore). The protein concentration was adjusted at 2 mg/mL. The reservoir solution consists of 50 mM sodium cacodylate-HCl pH 6.5 and 1.2 M ammonium sulfate. The crystals reached 0.2–0.4 mm within 1 week at 16°C. The crystals were transferred to a reservoir solution supplemented with 25% glycerol and flash-frozen at 100K. The X-ray diffraction experiments were carried out on beamlines of Photon Factory (Tsukuba) and of SPring-8 (Harima). The diffraction data used for the following structure determination were collected on BL17A of Photon Factory, and were processed with HKL2000 (***Otwinowski et al., 1997***). The phases were determined by molecular replacement with MOLREP (***Vagin et al., 2010***) using coordinates of CCD^WT^ in complex with the NCINI-related compound (PDB: 4GW6) as a search model (***Feng et al., 2013***). Structural refinements were carried out using PHENIX (***Adams et al., 2010***) and COOT (***Emsley et al., 2004***). Data collection and refinement statistics of CCD^P15^ (PDB: 6L0C) were listed in Figure 5—table supplement 1. Molecular graphics were prepared using PyMOL version 2.0 (Schrödinger, LLC).

### DSF

Recombinant CCD proteins (12.5, 25, and 50 µM) carrying A128T, H171Q, K173Q, and P15, respectively, together with F185K were prepared in sample buffer (50 mM HEPES pH 7.4, 500 mM NaCl, 7.5 mM CHAPS, 10 mM MgSO_4_, 5 mM BME, 5% Glycerol). SYPRO Orange (Life Technologies) was added to the samples (final concentration of SYPRO orange: 5×) (***Niesen et al., 2007***). The samples were successively heated from 25 to 95°C, and the increasing fluorescence intensity were detected by the real-time PCR system 7500 Fast (Applied Biosystems). Data was indicated as a relative ratio between minimum and maximum intensity of SYPRO orange from 25 to 95°C detected at each sample.

### SPR

SPR was examined using a Biacore X100 (GE Healthcare) at RT. Recombinant CCD proteins carrying A128T, K173Q, and A128T/K173Q mutations, respectively, together with F185K at 20 μg/mL diluted in HBS-EP buffer (GE Healthcare) were immobilized on two flow-cells of a CM5 sensor chip by direct amine coupling to 3,000 response units (RU). NCINI-3 diluted in HBS-EP buffer were flowed with a 2 min injection at 30 μL/min followed by a 2 min dissociation. KD values of NCINI-3 to the CCD proteins were calculated from the sensorgrams using the Biacore X100 control software.

### TEM

Viruses were produced by transfection with pNL_4-3_ IN^WT^ or IN^P26^ into 293T cells. The cells were washed at 12 h later, added fresh medium with or without 20 μM of NCINI-3. After 48 h incubation the samples were fixed in a solution of 4% formaldehyde and 4% glutaraldehyde in 0.1 M phosphate pH 7.4 buffer for 1 h at 4°C. After centrifugation, the supernatants were filtered through a 0.45 μm filter, and then performed by ultracentrifugation using KUBOTA model 7780 (KUBOTA, Japan) at 22,000 rpm for 5 h at 4°C. Virus pellets were resolved in minuscule quantity of 2% glutaraldehyde in 0.1 M phosphate pH 7.4 buffer and shipped to Tokai Electron Microscopy, Inc. Japan. The samples were dehydrated, infiltrated, and polymerized to a resin. Ultra-thin sections (50 nm) were stained with 2% uranyl acetate and observed under a TEM (JEM-1400Plus; JEOL Ltd., Tokyo, Japan). Digital imaged (3296×2472 pixels) were taken with a CCD camera (EM-14830RUBY2; JEOL Ltd., Tokyo, Japan). The virus images over 100 per the samples were classified under three types by the locations of RNP and the capsid core in the viral particles as previous reports (***Jurado et al., 2013***; ***Desimmie et al., 2013***).

### IN-RNA binding assay

Blunt ended viral DNA substrate (1-850 bp) containing 5’UTR region in pNL_4-3_ was obtained by PCR with following primers: (5’-GGT CTC TCT GGT TAG ACC AGA TCT GAG C-3’ and 5’-AAA CAT GGG TAT TAC TTC TGG GCT GAA AG-3’) using PrimeSTAR^®^HS (TAKARA) and purified by QIAquick Gel Extraction (QIAGEN). Single strand viral RNA was synthesized from the purified viral DNA substrate using ScriptMAX® Thermo T7 Transcription Kit (TOYOBO, Japan). After digested the template viral DNA, the viral RNA was purified by NucleoSpin® RNA Kit (TAKARA). Recombinant IN_His_ proteins (300 nM) was incubated with the synthetic viral RNA (100 and 300 nM) in assembly buffer (20 mM HEPES pH 7.4, 100 mM NaCl, 2 mM DTT, 7.5 mM MgCl_2_, 15 µM ZnCl_2_, 12% DMSO, 10% PEG 6000) for 3 h at 4°C. The IN_His_ proteins binding to the viral RNA were cross-linked with 0.05 mM BS3, and then visualized by immunoblotting.

### Alpha screening assay

Alpha screening 1 (***Demeulemeester et al., 2012***; ***Kessl et al., 2016***) was used the recombinant IN_His_ proteins (10 nM final concentration) which were incubated with various concentration of synthetic biotinylated TAR-RNA (biotin 5’-dT/GGU CUC UCU GGU UAG ACC AGA UCU GAG CCU GGG AGC UCU CUG GCU AAC UAG GGA ACC/3’-biotin dT) (Integrated DNA Technologies) at 4°C for 3 h in the AlphaLISA buffer (PerkinElmer). Anti-His Donor and Streptavidin Acceptor AlphaLISA beads (PerkinElmer) were added at final concentration 10 mg/mL to each well and incubated on RT for 3 h. The FRET signal was detected by an EnSpire Plate Reader (PerkinElmer) with an AlphaScreen module setting. Alpha screening 2 was used the recombinant IN_His_ and F-IN proteins which were incubated with various concentration of synthetic TAR-RNA (5’ GGU CUC UCU GGU UAG ACC AGA UCU GAG CCU GGG AGC UCU CUG GCU AAC UAG GGA ACC 3’) (Integrated DNA Technologies) at 4°C for 3 h in the AlphaLISA buffer. Anti-His Donor and anti-Flag Acceptor AlphaLISA beads were added at final concentration 10 mg/mL to each well and incubated on RT for 3 h, and then the FRET signal was also detected by the same above method.

### HIV-1 replication fitness

Wild type or recombinant HIV-1_NL4-3_ clones carrying E11K and NCINI-3 resistance mutations at 30 ng/ml of p24 were exposed to MT-4 cells (2.4 × 10^5^/ml) in TC 6-well plates for 3 h, and washed the cells, and divided into three fractions in fresh medium, and each cultured p24 levels were measured by HIV-1 p24 ELISA at 1, 3, 5, 7 days (***Amano et al., 2007***).

### Drug Susceptibility Assay

Evaluation for EC_50_ and CC_50_ of drugs and test compounds was previously reported (***Amano et al., 2007***). In brief, 3-(4,5-di-methylthiazol-2-yl)-2,5-diphenyltetrazolium bromide, yellow tetrazole (MTT) assay using MT-2 cells for determining EC_50_ represents inhibition of a 7 day multi-round HIV-1_LAI_ replication by detecting the living cells. Anti-HIV-1 assay (p24 assay) employing MT-4 cells represents inhibition of a 7-day multi-round HIV-1_NL4-3_ replication. EC_50_ was determined by measuring p24 levels using p24 ELISA. CC_50_s of the drugs and test compounds to MT-4 and HEK293T cells were determined using MTT assay.

### Compounds

NCINI-1, -2, -3, and DRV were synthesized using the published synthetic methods (***Christ et al., 2010***; ***Christ et al., 2012***; ***Ghosh et al., 2004***). RAL, EVG, and DTG were purchased from Selleck Chemicals, and BI224436 from Med Chem Express LLC.

## Acknowledgements

We also thank Haruo Aikawa and Hirokazu Tamamura for NCINIs synthesis and Sachiko Otsu for technical assistance of the experiments and Tokai Electron Microscopy Inc. staff for technical advice of TEM sample preparation and the data collection. We also thank the beamline staff at Photon Factory and SPring-8 for their help in the data collection. This work was supported by Japan Society for the Promotion of Science JP16K15521, JP18K16180, JP18K08436, and Development of Novel Drugs for Treating HIV-1 Infection and AIDS.

## Funding

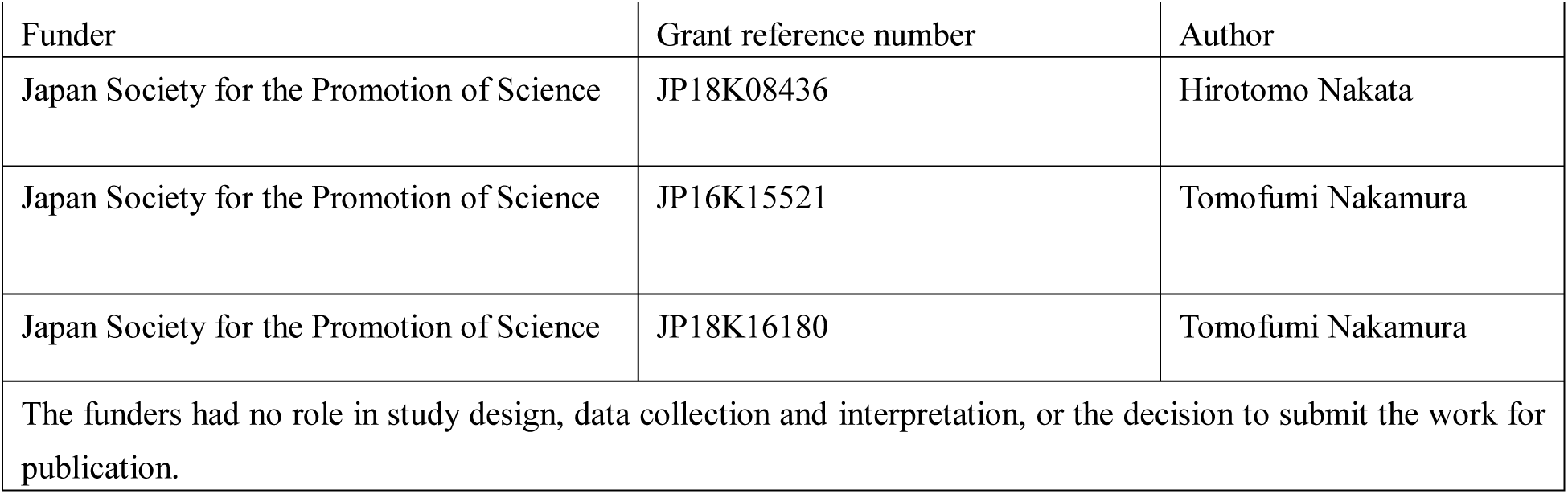

## Author contributions

**Tomofumi Nakamura**, Conceptualization, Methodology, Validation, Formal analysis, Investigation, Writing – original draft preparation, Visualization, Project administration, Funding acquisition; **Teruya Nakamura**, Methodology, Writing – original draft preparation, Software, Formal analysis, Investigation, Data curation, Visualization; **Masayuki Amano**, Methodology, Writing – original draft preparation; **Toshikazu Miyakawa**, Methodology, Writing – original draft preparation; **Yuriko Yamagata**, Methodology, Supervision; **Masao Matsuoka**, Writing – review & editing, Supervision, Project administration; **Hirotomo Nakata**, Conceptualization; Writing – original draft preparation; Writing – review & editing, Supervision; Project administration; Funding acquisition.

## Supplementary figures and tables

**Figure 1—figure supplement 1.**
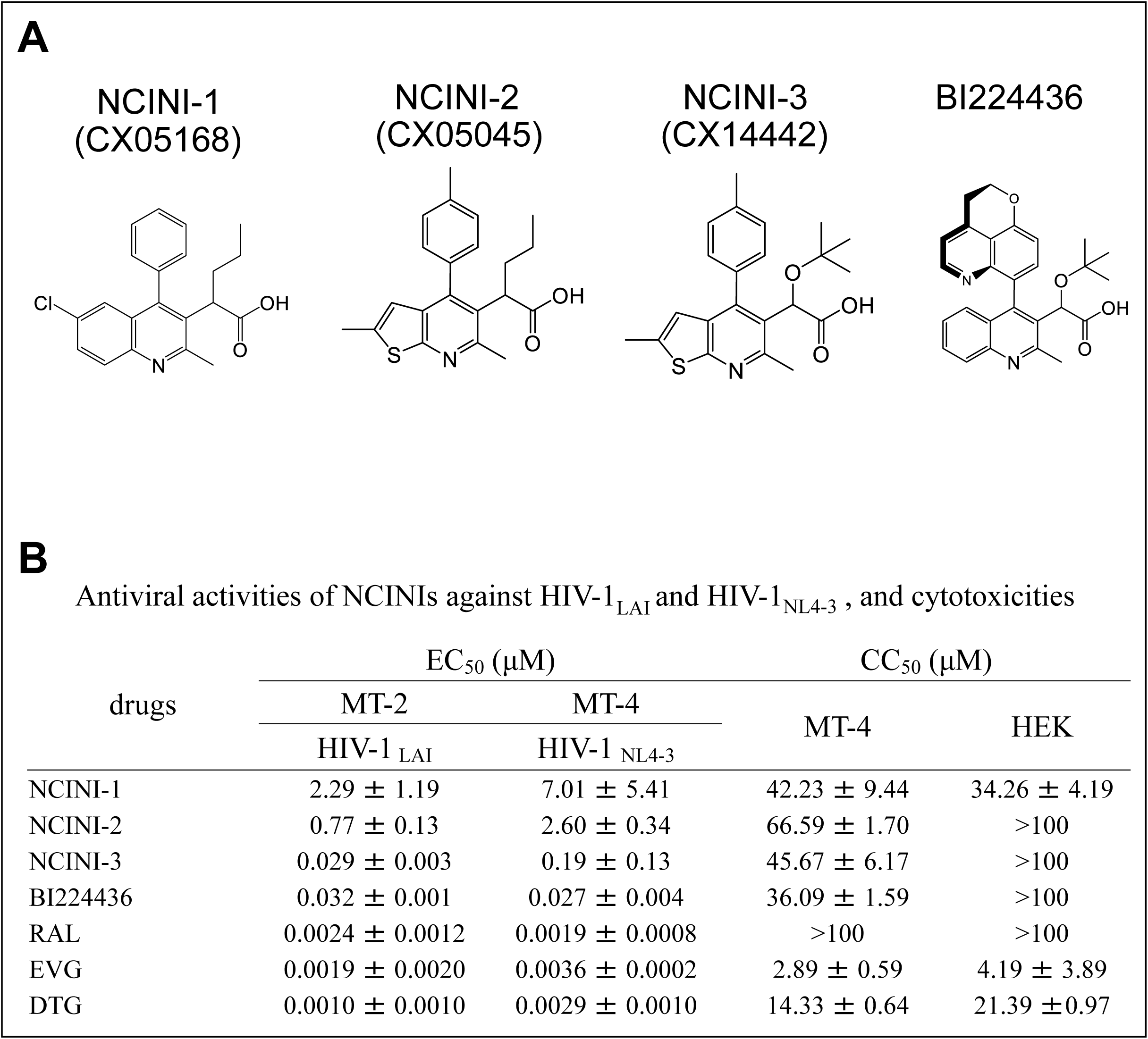
Compounds information. (**A**) Chemical structures of NCINIs. (**B**) Antiviral activities of NCINIs and INSTIs against HIV-1_LAI_ and HIV-1_NL4-3_, and cytotoxicities. MT-2 cells are exposed to 100 TCID_50_s of HIV-1_LAI_ and cultured in the presence of various concentration of NCINIs and INSTIs, and EC_50s_ are determined by using MTT assay. Cytotoxicities (CC_50s_) of NCINIs and INSTIs against MT-4 and HEK293T cells are also determined by MTT assay. All assays are conducted in duplicate, and data shown represent mean values (±SD) derived from results of three independent experiments.

**Figure 2—figure supplement 2.**
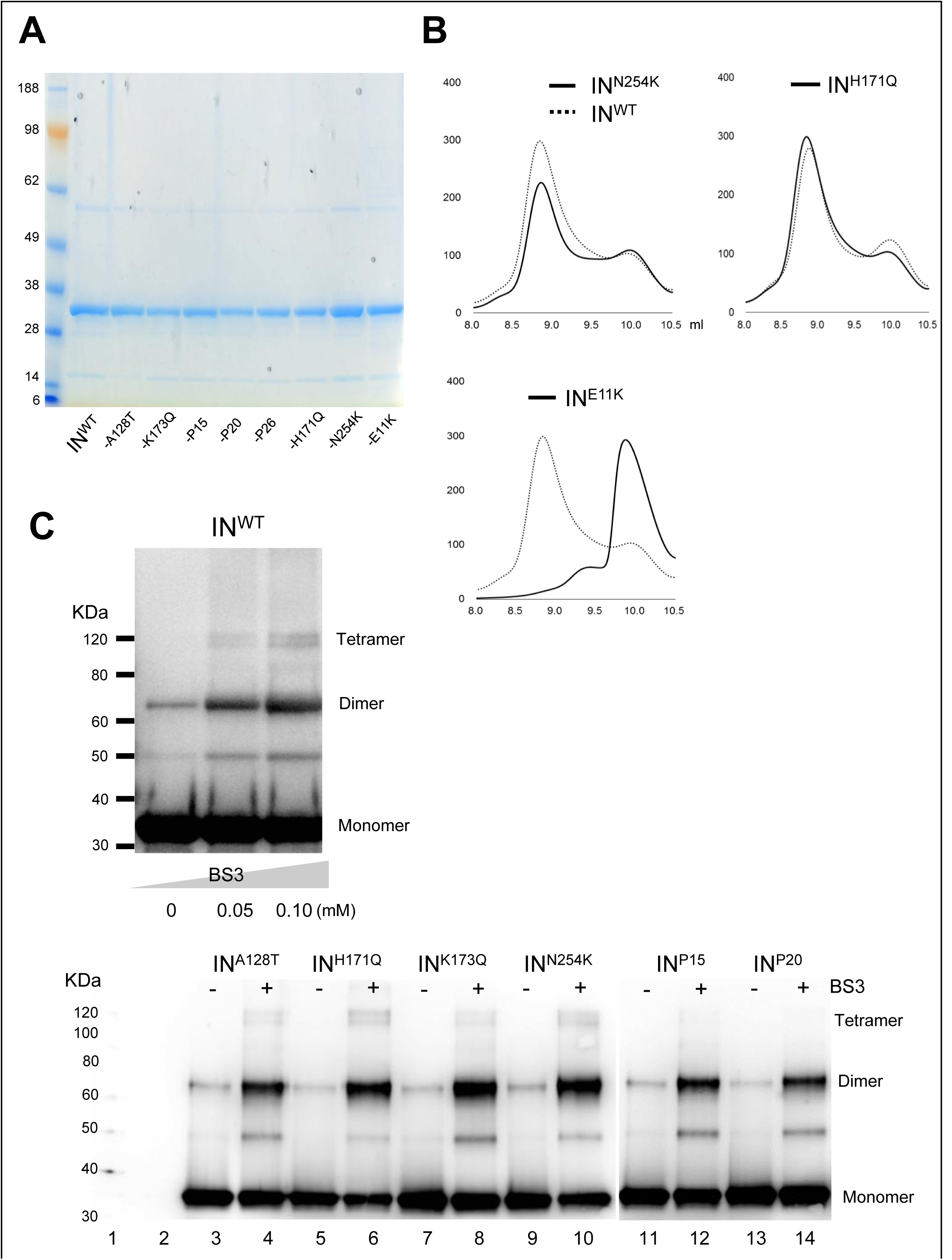
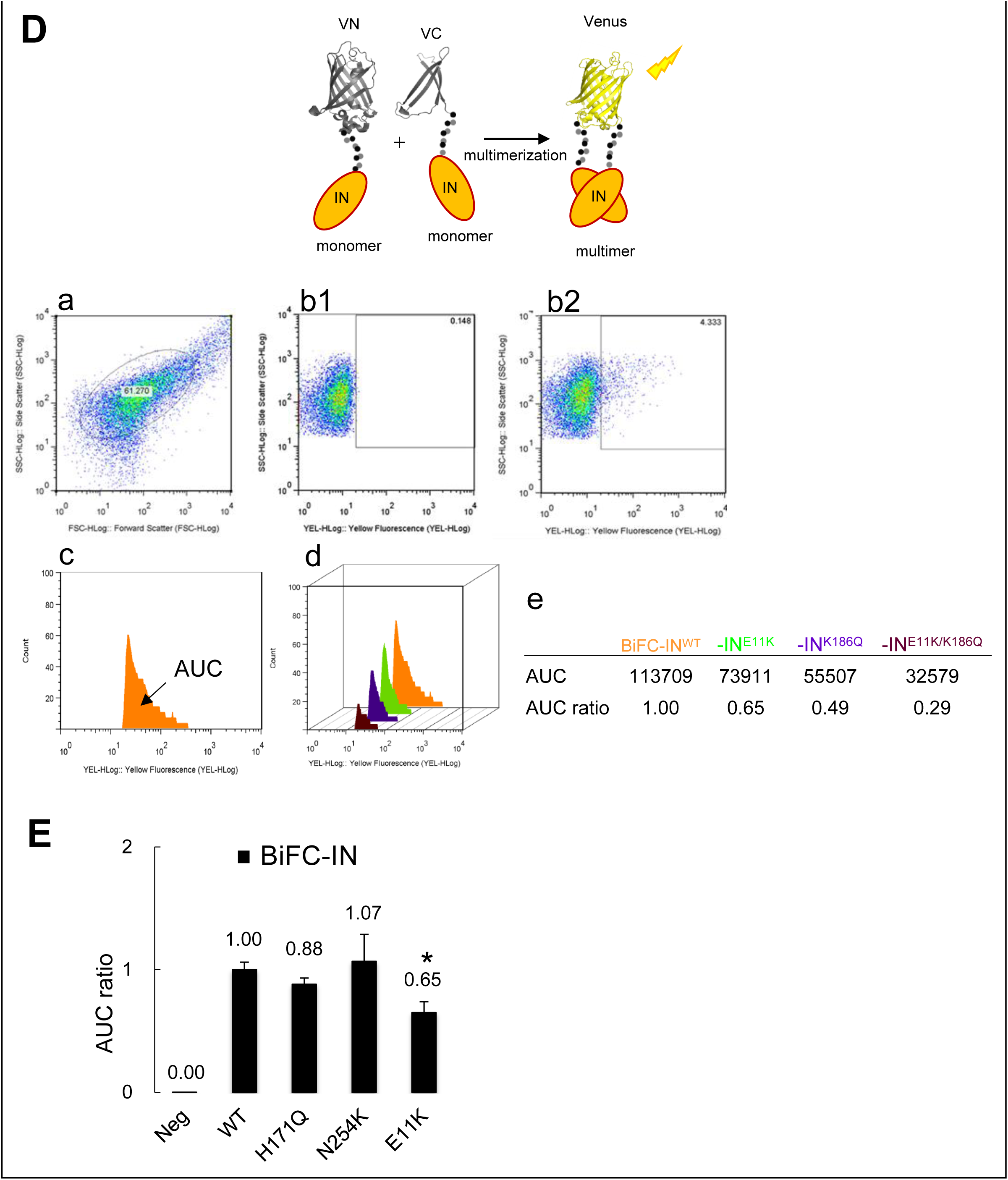
Purity and multimerization profile of IN_His_ proteins carrying NCINI-3 resistance mutations. (**A**) Purity of IN_His_ employed in this study. The recombinant proteins normalized with the proteins concentration are denatured and applied at 5 µg/well, and stained with Coomassie Brilliant Blue. (**B**) Multimerization profiles of IN^N254K^, IN^H171Q^, and IN^E11K^ using HPLC. (**C**) IN^WT^ proteins cross-linked with or without BS3 at 0.05 and 0.10 mM, respectively, are visualized by immunoblotting with anti-IN antibody at upper panel. IN^A128T^, IN^H171Q^, IN^K173Q^, IN^N254K^, IN^P15^, and IN^P20^ cross-linked with (Lanes 4, 6, 8, 10, 12, and 14) or without (lanes 3, 5, 7, 9, 11, and 13) BS3 at 0.05 mM are visualized by immunoblotting with anti-IN antibody at lower panel. (**D**) Schematic illustration of BiFC-IN system is shown at upper panel. BiFC-IN system procedure using FACS at middle panel. HEK293T cells are gated with SSC and FSC (a). Negative control (b1) and BiFC-IN^WT^ data (b2) selected from Gate a. Area under the curve (AUC) of BiFC-IN^WT^ which comprises Venus intensity and cell count of Venus positive cells selected from Gate b2 (c). AUCs of BiFC-IN^WT^ and -IN^E11K^, -IN^K186Q^, and -IN^E11K/K186Q^ (d) are measured by ImageJ, and AUC ratios of BiFC-IN^E11K^, -IN^K186Q^, and -IN^E11K/K186Q^ to BiFC-IN^WT^ are shown (e). These mutations are known to decrease the IN multimerization. (**E**) AUC ratios of BiFC-IN^WT^, -IN^H171Q^, -IN^N254K^, and -IN^E11K^, respectively are shown. The AUC ratios are mean values (±SD) derived from FACS results of three independent experiments. Statistical significance was examined using two-tailed Student t-test, * P < 0.05.

**Figure 2—table supplement 1.**
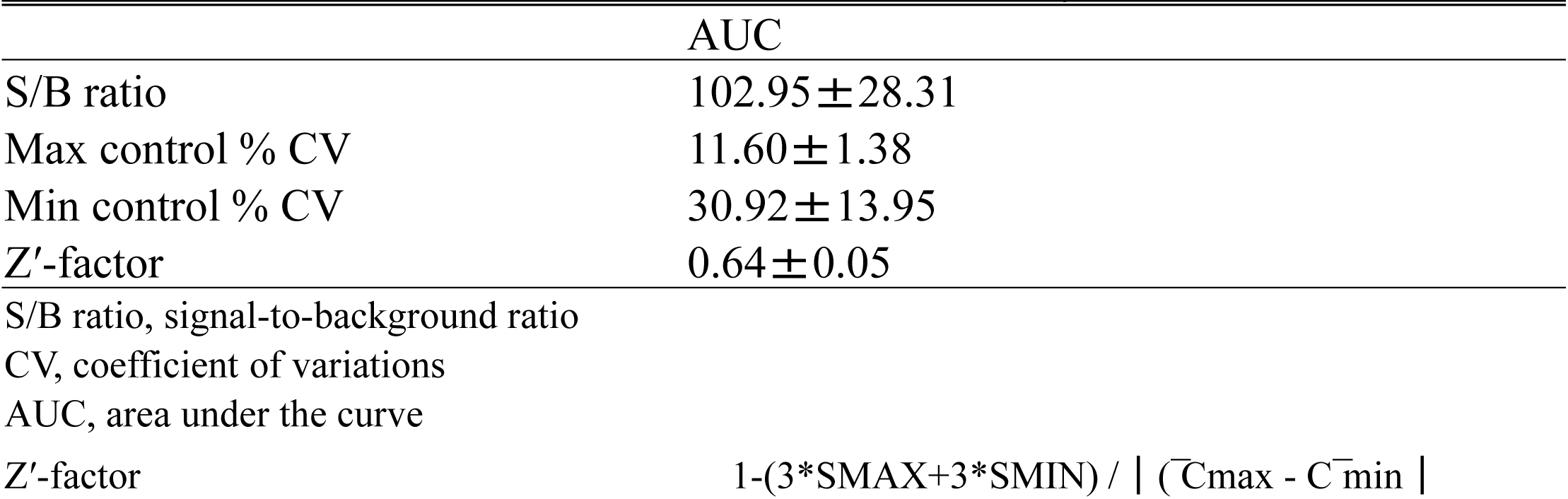
Statistical evaluation of BiFC-IN system. Statistical indices in BiFC-IN system are reported previously (***Nakamura et al., 2016***).

**Figure 3—figure supplement 3.**
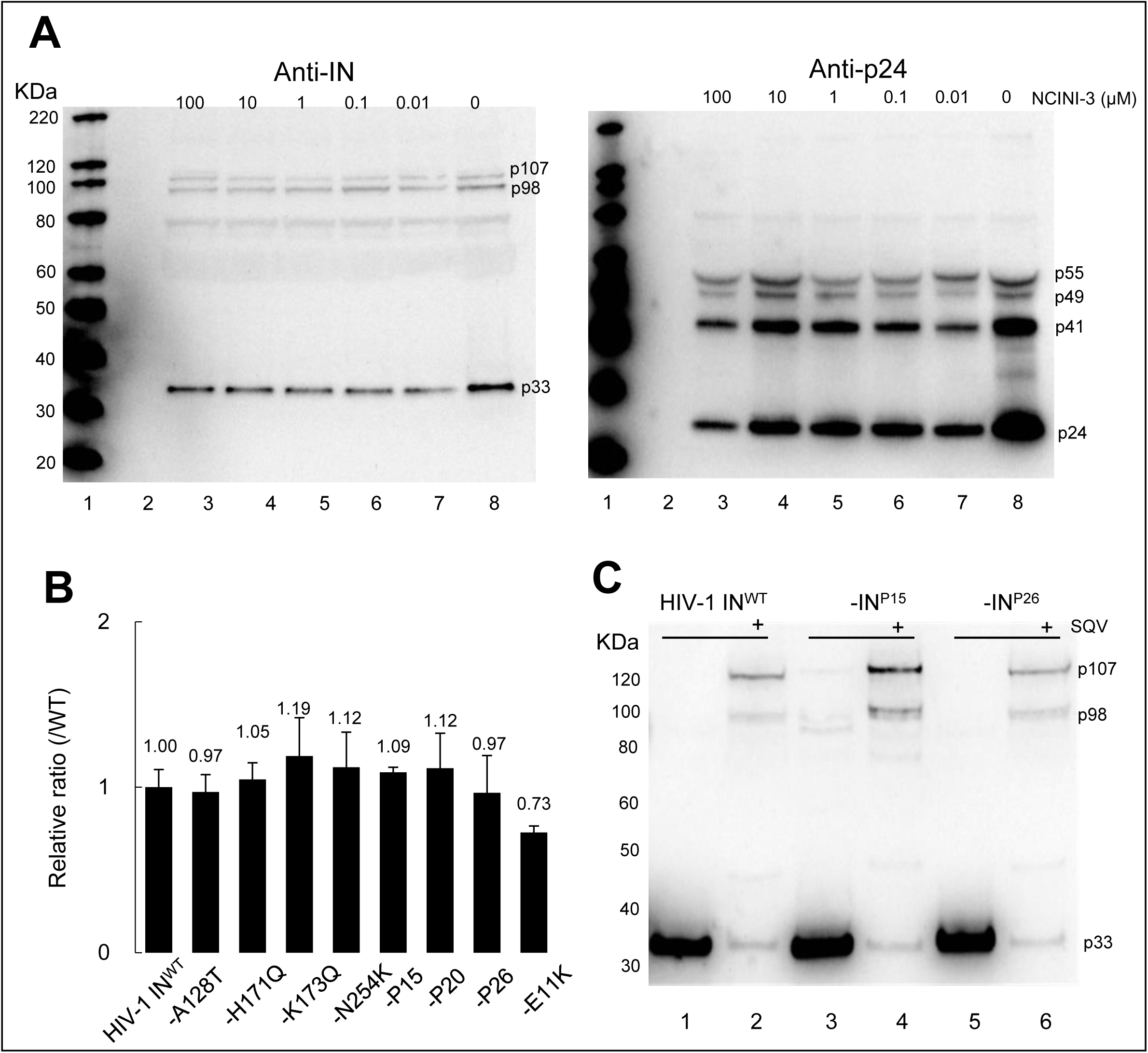
Effect of NCINIs on Gag and Gag-pol proteolytic processing and viral production of recombinant HIV-1_NL4-3_ clones carrying the mutations used in this study. (**A**) Gag and Gag-pol proteolytic processing of crude HIV-1 IN^WT^ viral particles. The samples are generated by transfection experiments in the various concentration from 0.01 to 100 μM of NCINI-3, and visualized by SDS PAGE with anti-IN antibody at left and anti-p24 antibody at right panel. The representative results are shown from two independent experiments. (**B**) Viral production, p24 levels of HIV-1 IN^WT^ and HIV-1 clones carrying NCINI-3 resistance mutations and E11K mutation. Data are normalized to HIV-1 IN^WT^ and represent mean values (±SD) derived from three independent experiments. (**C**) Pol proteolytic processing of purified HIV-1 IN^WT^, -IN^P15^, and, -IN^P26^ viral particles generated in the presence (Lanes 2, 4, and, 6) or absence (Lanes 1, 3, and 5) of 2 μM SQV are visualized by immunoblotting with anti-IN antibody. All data are from two independent experiments and the representative results are shown.

**Figure 3—table supplement 1.** Replication kinetics of wild type and recombinant HIV-1 clones propagating within MT-4 cells. All assays are conducted in duplicate, and data shown represent mean values (±SD) derived from results of three independent experiments.

| recombinant HIV-1<sub>NL4-3</sub> clones | p24 (ng/ml) |  |  |  |
| --- | --- | --- | --- | --- |
|  | Day 1 | 3 | 5 | 7 |
| HIV-1<sub>NL4-3</sub> IN<sup>WT</sup> | 0.0 | 30.4 ± 34.7 | 830.3 ± 198.4 | 1312.5 ± 57.7 |
| -IN<sup>A128T</sup> | 0.0 | 49.1 ± 18.0 | 674.6 ± 102.3 | 1030.6 ± 118.0 |
| -IN<sup>H171Q</sup> | 0.0 | 71.1 ± 40.1 | 945.7 ± 126.5 | 1220.9 ± 192.6 |
| -IN<sup>K173Q</sup> | 0.0 | 21.6 ± 23.6 | 1073.6 ± 283.4 | 1671.9 ± 461.0 |
| -IN<sup>N254K</sup> | 0.0 | 18.7 ± 21.8 | 979.7 ± 233.4 | 1601.4 ± 229.5 |
| -IN<sup>P15</sup> | 0.0 | 42.2 ± 11.5 | 970.6 ± 81.1 | 1306.5 ± 265.0 |
| -IN<sup>P20</sup> | 0.0 | 24.7 ± 19.1 | 1034.5 ± 63.3 | 1412.2 ± 215.8 |
| -IN<sup>P26</sup> | 0.0 | 10.1 ± 14.2 | 794.9 ± 352.2 | 1443.1 ± 340.0 |
| -IN<sup>E11K</sup> | 0.0 | 0.0 | 0.0 | 0.0 |

**Figure 4—figure supplement 4.**
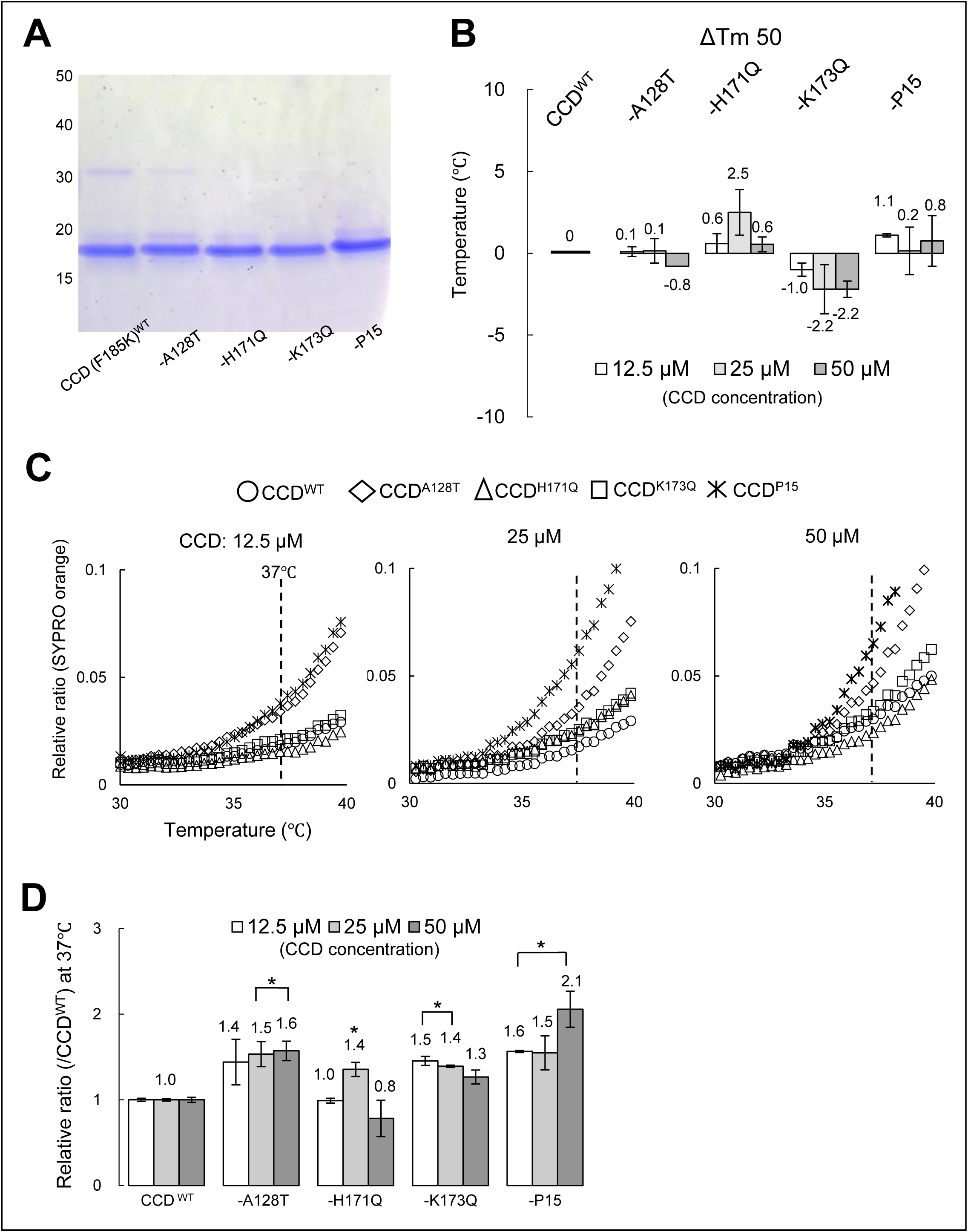
Purity and stability of CCD_His_ carrying NCINI-3 resistance mutation together with F185K mutation. (**A**) Purity of the CCD employed in this study. The recombinant CCD are denatured and applied at 5 µg/well, and stained with Coomassie Brilliant Blue. (**B**) ΔTm 50 of the CCD proteins by DSF. ΔTm 50 indicates Tm 50 differences between CCD^WT^ and each CCD carrying the mutations at 12.5, 25, and 50 μM concentration, respectively. (**C**) Relative ratios of SYPRO orange of CCD^WT^, CCD^A128T^, CCD^H171Q^, CCD^K173Q^, and CCD^P15^ from 30 to 40°C are shown at 12.5, 25, and 50 μM concentration, respectively. (**D**) Proportion of surface hydrophobic sites of CCD proteins at 37°C. Relative ratios to CCD^WT^ obtained from SYPRO orange binding to the surface hydrophobic sites of CCD^A128T^, CCD^H171Q^, CCD^K173Q^, and CCD^P15^, respectively are shown at 12.5, 25, and 50 μM concentration, respectively. Data represents mean values (±SD) derived from three independent experiments. Statistical significance is examined using two-tailed Student t-test, * P < 0.05.

**Figure 5—figure supplement 5.**
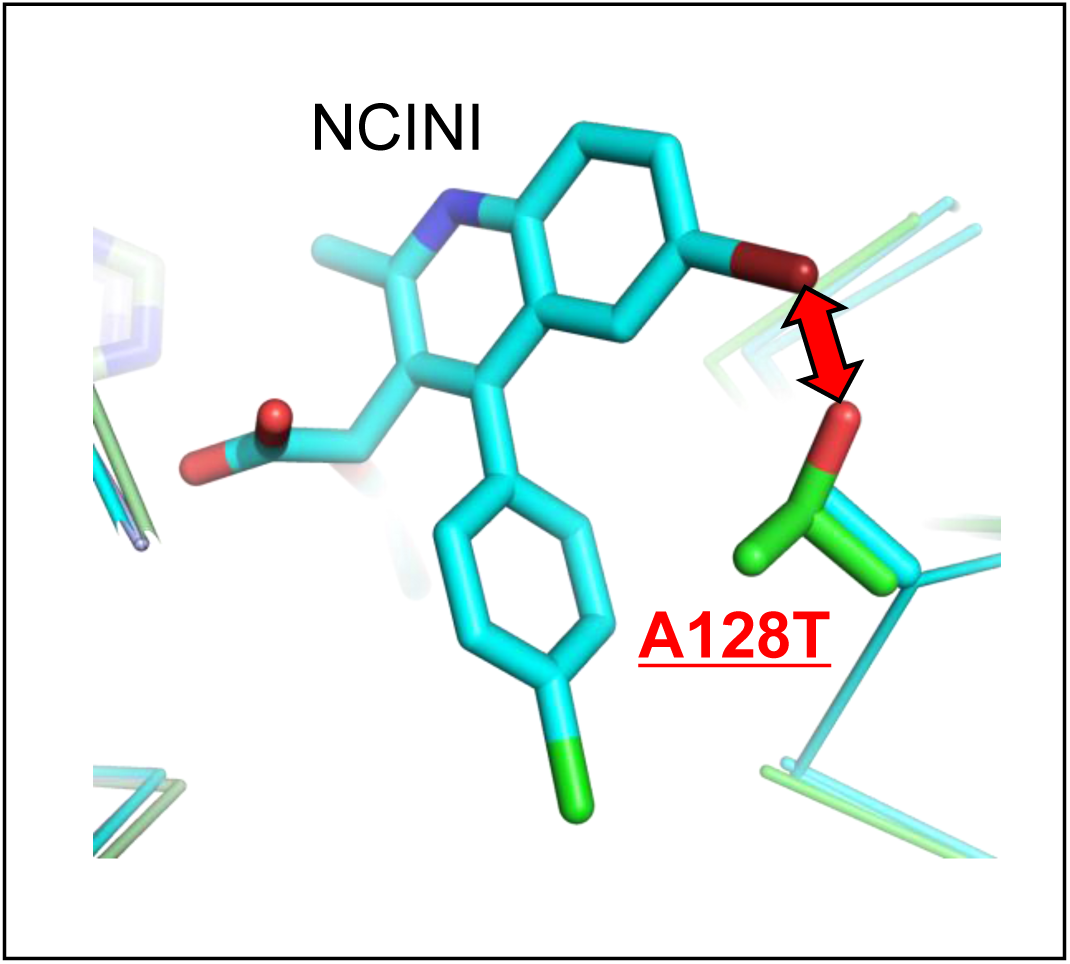
Effect of A128T mutation on the position of the NCINI-related compound. The NCINI is shifted to the outside of the NCINI binding pocket by the side chain of T128 residue. Overlay of the crystal structures of the NCINI-related compound binding to CCD^WT^ (PDB: 4GW6) and CCD^P15^ (PDB: 6L0C), respectively.

**Figure 5—table supplement 1.**
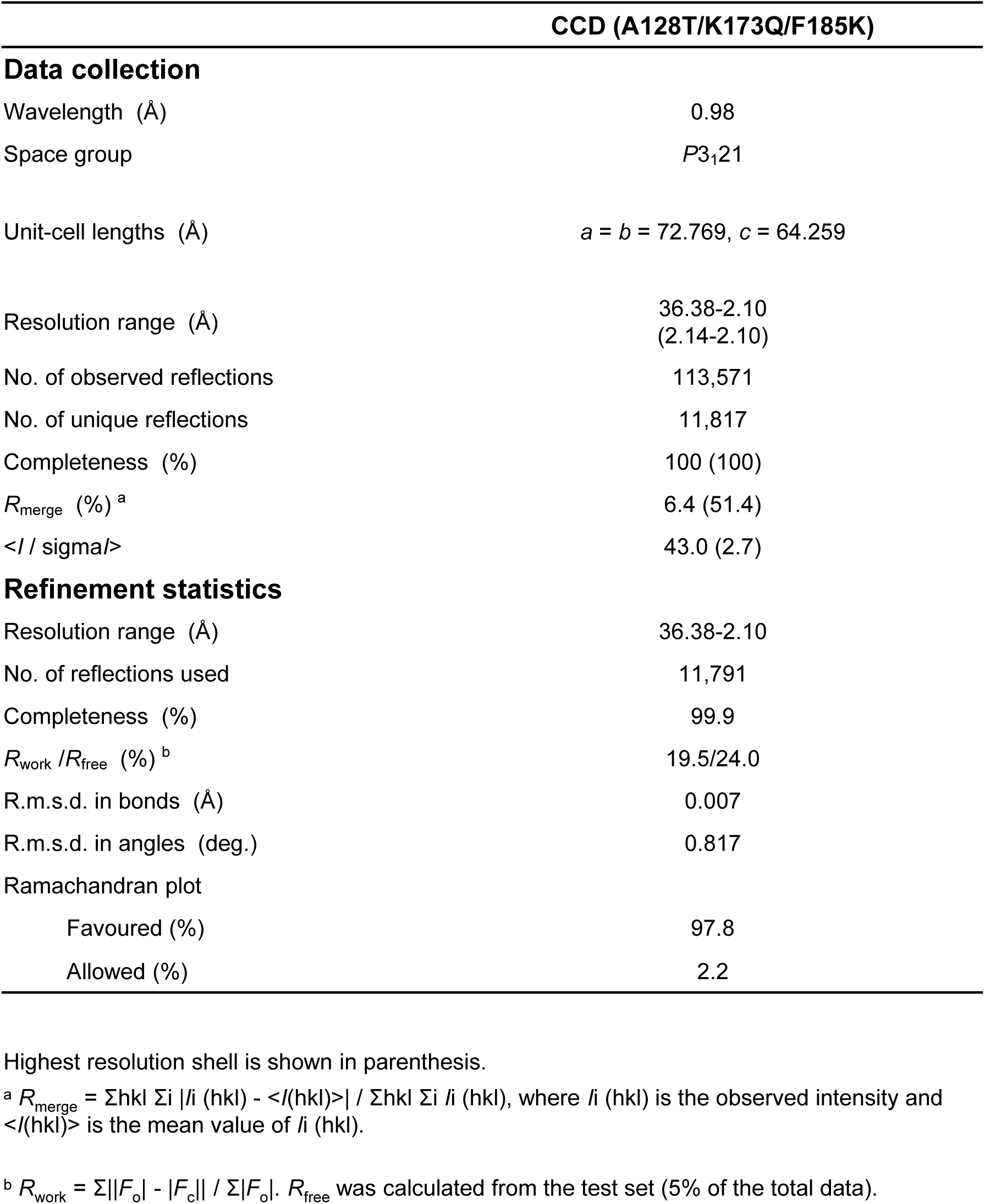
X-ray crystal structure data collection and refinements.

**Figure 6—figure supplement 6.**
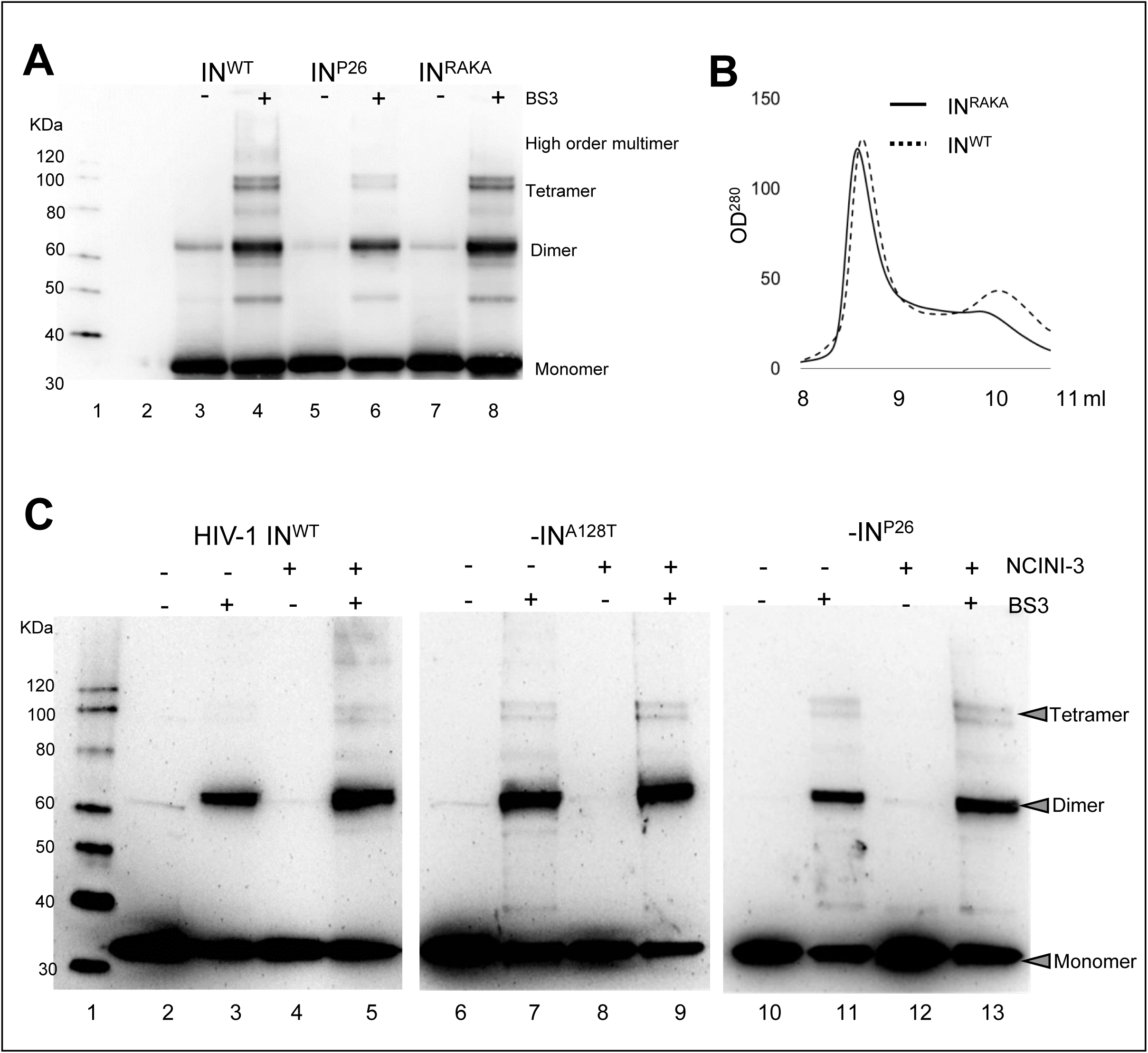
Multimerization profile of IN proteins. (**A**) Multimerization ratio of purified IN proteins (IN^WT^, IN^P26^, and IN^RAKA^) cross-linking with BS3. MW (Lane 1), buffer (Lane 2), IN^WT^, IN^P26^, and IN^RAKA^ (Lanes 3, 5, and 7), and IN^WT^, IN^P26^, and IN^RAKA^ cross-linked with 0.05 mM BS3 (Lanes 4, 6, and 8) are shown. (**B**) Multimerization profiles of IN^WT^ and IN^RAKA^ using HPLC. (**C**) IN multimerization in HIV-1_NL4-3_ IN^WT^, -IN^A128T^, and -IN^P26^ viral particles generated in the presence or absence of 20 μM NCINI-3. The HIV-1 IN^WT^, -IN^A128T^, and -IN^P26^ clones are purified by ultracentrifugation. MW (Lane 1), the high concentrated HIV-1 IN^WT^, - IN^A128T^, and -IN^P26^ stripped the viral membrane by 0.5% triton X PBS are cross-linked with (Lanes 3, 5, 7, 9, 11, and 13) or without (Lanes 2, 4, 6, 8, 10, and 12) BS3 and employed 100 ng of p24 at each sample per wells, and visualized by SDS-PAGE with anti-HIV-1 IN antibody.

## References

Adams PD, Afonine PV, Bunkóczi G, Chen VB, Davis IW, Echols N, Headd JJ, Hung LW, Kapral GJ, Grosse-Kunstleve RW, McCoy AJ, Moriarty NW, Oeffner R, Read RJ, Richardson DC, Richardson JS, Terwilliger TC, Zwart PH. 2010. PHENIX : a comprehensive Python-based system for macromolecular structure solution. Acta Crystallogr. Sect. D Biol. Crystallogr. 66:213–221. doi.10.1107/S0907444909042589.

Amano M, Koh Y, Das D, Li J, Leschenko S, Wang YF, Boross PI, Weber IT, Ghosh AK, Mitsuya H. 2007. A novel bis-tetrahydrofuranylurethane-containing nonpeptidic protease inhibitor (PI), GRL-98065, is potent against multiple-PI-resistant human immunodeficiency virus in vitro. Antimicrob Agents Chemother 51:2143–55. doi:10.1128/AAC.01413-06.

Balakrishnan M, Yant SR, Tsai L, O Sullivan C, Bam RA, Tsai A, Niedziela-Majka A, Stray KM, Sakowicz R, Cihlar T. 2013. Non-Catalytic Site HIV-1 Integrase Inhibitors Disrupt Core Maturation and Induce a Reverse Transcription Block in Target Cells. PLoS ONE 8:e74163. doi:10.1371/journal.pone.0074163.

Cherepanov P, Maertens G, Proost P, Devreese B, Van Beeumen J, Engelborghs Y, De Clercq E, Debyser Z. 2003. HIV-1 integrase forms stable tetramers and associates with LEDGF/p75 protein in human cells. J Biol Chem 278:372–81. doi:10.1074/jbc.M209278200.

Christ F, Voet A, Marchand A, Nicolet S, Desimmie BA, Marchand D, Bardiot D, der Veken N, Remoortel B, Strelkov SV, Maeyer M, Chaltin P, Debyser Z. 2010. Rational design of small-molecule inhibitors of the LEDGF/p75-integrase interaction and HIV replication. Nature Chemical Biology 6:442–448. doi:10.1038/nchembio.370.

Christ F, Shaw S, Demeulemeester J, Desimmie BA, Marchand A, Butler S, Smets W, Chaltin P, Westby M, Debyser Z, Pickford C. 2012. Small-molecule inhibitors of the LEDGF/p75 binding site of integrase block HIV replication and modulate integrase multimerization. Antimicrob Agents Chemother 56:4365–74. doi:10.1128/AAC.00717-12.

Demeulemeester J, Tintori C, Botta M, Debyser Z, Christ F. 2012. Development of an AlphaScreen-based HIV-1 integrase dimerization assay for discovery of novel allosteric inhibitors. J Biomol Screen 17:618–28. doi:10.1177/1087057111436343.

Desimmie BA, Schrijvers R, Demeulemeester J, Borrenberghs D, Weydert C, Thys W, Vets S, Remoortel VB, Hofkens J, Rijck DJ, Hendrix J, Bannert N, Gijsbers R, Christ F, Debyser Z. 2013. LEDGINs inhibit late stage HIV-1 replication by modulating integrase multimerization in the virions. Retrovirology 10:57. doi:10.1186/1742-4690-10-57.

Dowers E, Zamora F, Barakat LA, Ogbuagu O. 2018. Dolutegravir/rilpivirine for the treatment of HIV-1 infection. HIV AIDS (Auckl) 10:215–224. doi:10.2147/HIV.S157855.

Dyda F, Hickman AB, Jenkins TM, Engelman A, Craigie R, Davies DR. 1994. Crystal structure of the catalytic domain of HIV-1 integrase: similarity to other polynucleotidyl transferases. Science 266:1981–6. doi:10.1126/science.7801124.

Emsley P, Cowtan K. 2004. Coot: model-building tools for molecular graphics. Acta Crystallogr. Sect. D Biol. Crystallogr. 60:2126–32. doi:10.1107/S0907444904019158.

Engelman A. 1999. In vivo analysis of retroviral integrase structure and function. Adv Virus Res. 52:411–26. doi:10.1016/S0065-3527.60309-7.

Feng L, Sharma A, Slaughter A, Jena N, Koh Y, Shkriabai N, Larue RC, Patel PA, Mitsuya H, Kessl JJ, Engelman A, Fuchs JR, Kvaratskhelia M. 2013. The A128T resistance mutation reveals aberrant protein multimerization as the primary mechanism of action of allosteric HIV-1 integrase inhibitors. J Biol Chem 288:15813–20. doi:10.1074/jbc.M112.443390.

Fenwick C, Amad M, Bailey MD, Bethell R, Bös M, Bonneau P, Cordingley M, Coulombe R, Duan J, Edwards P, Fader LD, Faucher A-MM, Garneau M, Jakalian A, Kawai S, Lamorte L, LaPlante S, Luo L, Mason S, Poupart M-AA, Rioux N, Schroeder P, Simoneau B, Tremblay S, Tsantrizos Y, Witvrouw M, Yoakim C. 2014. Preclinical profile of BI 224436, a novel HIV-1 non-catalytic-site integrase inhibitor. Antimicrob Agents Chemother 58:3233–44. doi:10.1128/AAC.02719-13.

Flexner C, Tierney C, Gross R, Andrade A, Lalama C, Eshleman S, Aberg J, Sanne I, Parsons T, Kashuba A, Rosenkranz S, Kmack A, Ferguson E, Dehlinger M, Mildvan D. 2010. Comparison of Once-Daily versus Twice-Daily Combination Antiretroviral Therapy in Treatment-Naive Patients: Results of AIDS Clinical Trials Group (ACTG) A5073, a 48-Week Randomized Controlled Trial. Clin Infect Dis 50:1041–1052. doi:10.1086/651118.

Fontana J, Jurado KA, Cheng N, Ly NL, Fuchs JR, Gorelick RJ, Engelman AN, Steven AC. 2015. Distribution and Redistribution of HIV-1 Nucleocapsid Protein in Immature, Mature, and Integrase-Inhibited Virions: a Role for Integrase in Maturation. J Virol 89:9765–80. doi:10.1128/JVI.01522-15

Ghosh AK, Leshchenko S, Noetzel M. 2004. Stereoselective photochemical 1,3-dioxolane addition to 5-alkoxymethyl-2(5H)-furanone: synthesis of bis-tetrahydrofuranyl ligand for HIV protease inhibitor UIC-94017 (TMC-114). J Org Chem 69:7822–9. doi:10.1021/jo049156y

Gulick RM, Flexner C. 2019. Long-Acting HIV Drugs for Treatment and Prevention. Annu Rev Med 70:137–150. doi:10.1146/annurev-med-041217-013717.

Hoyte AC, Jamin AV, Koneru PC, Kobe MJ, Larue RC, Fuchs JR, Engelman AN, Kvaratskhelia M. 2017. Resistance to pyridine-based inhibitor KF116 reveals an unexpected role of integrase in HIV-1 Gag-Pol polyprotein proteolytic processing. J Biol Chem 292:19814–19825. doi:10.1074/jbc.M117.816645.

Hare S, Nunzio DF, Labeja A, Wang J, Engelman A, Cherepanov P. 2009. Structural basis for functional tetramerization of lentiviral integrase. PLoS Pathog 5:e1000515. doi:10.1371/journal.ppat.1000515.

Hare S, Gupta SS, Valkov E, Engelman A, Cherepanov P. 2010. Retroviral intasome assembly and inhibition of DNA strand transfer. Nature 464:232–6. doi:10.1038/nature0878.

Huldrych F Günthard Vincent Calvez Roger Paredes Deenan Pillay Robert W Shafer Annemarie M Wensing Donna M Jacobsen Douglas D Richman, Human Immunodeficiency Virus Drug Resistance: 2018 Recommendations of the International Antiviral Society–USA Panel, 2019, Clinical Infectious Diseases, 68 (2) 177–187, https://doi.org/10.1093/cid/ciy463.

Jurado KA, Wang H, Slaughter A, Feng L, Kessl JJ, Koh Y, Wang W, Ballandras-Colas A, Patel PA, Fuchs JR, Kvaratskhelia M, Engelman A. 2013. Allosteric integrase inhibitor potency is determined through the inhibition of HIV-1 particle maturation. Proceedings of the National Academy of Sciences 110:8690–8695. doi:10.1073/pnas.1300703110.

Kanters S, Park JJ, Chan K, Socias ME, Ford N, Forrest JI, Thorlund K, Nachega JB, Mills EJ. 2017. Interventions to improve adherence to antiretroviral therapy: a systematic review and network meta-analysis. Lancet HIV 4:e31–e40. doi:10.1016/S2352-3018(16)30206-5.

Kessl JJ, Jena N, Koh Y, Taskent-Sezgin H, Slaughter A, Feng L, de Silva S, Wu L, Le Grice SF, Engelman A, Fuchs JR, Kvaratskhelia M. 2012. Multimode, cooperative mechanism of action of allosteric HIV-1 integrase inhibitors. J Biol Chem 287:16801–11. doi:10.1074/jbc.M112.354373.

Kessl JJ, Kutluay SB, Townsend D, Rebensburg S, Slaughter A, Larue RC, Shkriabai N, Bakouche N, Fuchs JR, Bieniasz PD, Kvaratskhelia M. 2016. HIV-1 Integrase Binds the Viral RNA Genome and Is Essential during Virion Morphogenesis. Cell 166:1257–1268.e12. doi:10.1016/j.cell.2016.07.044

Lu R, Limón A, Devroe E, Silver PA, Cherepanov P, Engelman A. 2004. Class II integrase mutants with changes in putative nuclear localization signals are primarily blocked at a postnuclear entry step of human immunodeficiency virus type 1 replication. J Virol 78:12735–46. doi:10.1128/JVI.78.23.12735-12746.2004.

May MT, Gompels M, Delpech V, Porter K, Orkin C, Kegg S, Hay P, Johnson M, Palfreeman A, Gilson R, Chadwick D, Martin F, Hill T, Walsh J, Post F, Fisher M, Ainsworth J, Jose S, Leen C, Nelson M, Anderson J, Sabin C. 2014. Impact on life expectancy of HIV-1 positive individuals of CD4+ cell count and viral load response to antiretroviral therapy. AIDS 28:1193–202. doi:10.1097/QAD.0000000000000243.

Nakamura T, Campbell JR, Moore AR, Otsu S, Aikawa H, Tamamura H, Mitsuya H. 2016. Development and validation of a cell-based assay system to assess human immunodeficiency virus type 1 integrase multimerization. J Virol Methods 236:196–206. doi:10.1016/j.jviromet.2016.07.023.

Niesen FH, Berglund H, Vedadi M. 2007. The use of differential scanning fluorimetry to detect ligand interactions that promote protein stability. Nat Protoc 2:2212–2221. doi:10.1038/nprot.2007.321.

Otwinowski Z, Minor W. 1997. Processing of X-ray diffraction data collected in oscillation mode. Methods Enzymol. 276:307–26. doi:10.1016/S0076-6879.76066-X.

Passos DO, Li M, Yang R, Rebensburg SV, Ghirlando R, Jeon Y, Shkriabai N, Kvaratskhelia M, Craigie R, Lyumkis D. 2017. Cryo-EM structures and atomic model of the HIV-1 strand transfer complex intasome. Science 355:89–92. doi:10.1126/science.aah5163.

Rouzic LE, Bonnard D, Chasset S, Bruneau JM, Chevreuil F, Strat LF, Nguyen J, Beauvoir R, Amadori C, Brias J, Vomscheid S, Eiler S, Levy N, Delelis O, Deprez E, Saib A, Zamborlini A, Emiliani S, Ruff M, Ledoussal B, Moreau F, Benarous R. 2013. Dual inhibition of HIV-1 replication by integrase-LEDGF allosteric inhibitors is predominant at the post-integration stage. Retrovirology 10:144. doi:10.1186/1742-4690-10-144.

Sharma A, Slaughter A, Jena N, Feng L, Kessl JJ, Fadel HJ, Malani N, Male F, Wu L, Poeschla E, Bushman FD, Fuchs JR, Kvaratskhelia M. 2014. A new class of multimerization selective inhibitors of HIV-1 integrase. PLoS Pathog 10:e1004171. doi:10.1371/journal.ppat.1004171.

Slaughter A, Jurado K, Deng N, Feng L, Kessl J, Shkriabai N, Larue R, Fadel H, Patel P, Jena N, Fuchs J, Poeschla E, Levy R, Engelman A, Kvaratskhelia M. 2014. The mechanism of H171T resistance reveals the importance of Nδ-protonated His171 for the binding of allosteric inhibitor BI-D to HIV-1 integrase. Retrovirology 11:100. doi:10.1186/s12977-014-0100-1.

Trickey A, May MT, Vehreschild JJ, Obel N, Gill MJ, Crane HM, Boesecke C, Patterson S, Grabar S, Cazanave C, Cavassini M, Shepherd L, Monforte AD, van Sighem A, Saag M, Lampe F, Hernando V, Montero M, Zangerle R, Justice AC, Sterling T, Ingle SM, Sterne JAC. 2017. Antiretroviral Therapy Cohort Collaboration, Survival of HIV-positive patients starting antiretroviral therapy between 1996 and 2013: a collaborative analysis of cohort studies, Lancet HIV., 4 (8) e349–e356. doi: 10.1016/S2352-3018(17)30066-8.

Tsiang M, Jones GS, Niedziela-Majka A, Kan E, Lansdon EB, Huang W, Hung M, Samuel D, Novikov N, Xu Y, Mitchell M, Guo H, Babaoglu K, Liu X, Geleziunas R, Sakowicz R. 2012. New class of HIV-1 integrase (IN) inhibitors with a dual mode of action. J Biol Chem 287:21189–203. doi:10.1074/jbc.M112.347534.

Vagin A, Teplyakov A. 2010. Molecular replacement with MOLREP. Acta Crystallogr. Sect. D Biol. Crystallogr. 66:22–5. doi:10.1107/S0907444909042589.

